# Single-cell RNA-seq reveals a concomitant delay in differentiation and cell cycle of aged hematopoietic stem cell

**DOI:** 10.1101/2020.06.17.156893

**Authors:** Léonard Hérault, Mathilde Poplineau, Adrien Mazuel, Nadine Platet, Élisabeth Remy, Estelle Duprez

**Affiliations:** Epigenetic Factors in Normal and Malignant Hematopoiesis Team, Aix Marseille Université, CNRS, INSERM, Institut Paoli-Calmettes, CRCM, Marseille, France; Aix Marseille Université, CNRS, Centrale Marseille, I2M, Marseille, France

## Abstract

Hematopoietic stem cells (HSCs) are the guarantor of the proper functioning of hematopoiesis due to their incredible diversity of potential. During aging the heterogeneity of mouse HSCs evolves, which contributes to the deterioration of the immune system. Here we address the transcriptional plasticity of HSC upon aging at the single-cell resolution. Through the analysis of 15,000 young and aged transcriptomes, we reveal 15 clusters of HSCs unveiling rare and specific HSC abilities that change with age. Pseudotime ordering complemented with regulon analysis showed that the consecutive differentiation states of HSC are delayed upon aging. By analysing cell cycle at the single cell level we highlight an imbalance of cell cycle regulators of very immature aged HSC that may contribute to their accumulation in an undifferentiated state.

Our results therefore establish a reference map of young and old mouse HSC differentiation and reveal a potential mechanism that delay aged HSC differentiation.

## INTRODUCTION

The hematopoietic stem cell (HSC) is an adult tissue stem cell residing in the bone marrow (BM), with multipotent differentiation, regenerative and self-renewal abilities, the proper functioning of which is a guarantee of a healthy immune system. HSC properties have been extensively studied thanks to the use of specific surface markers and multicolored fluorescence-assisted cell sorting (FACS) analyses that have made it possible to isolate them and test their properties during serial grafts ^1, 2^. This cell-surface marker-based HSC characterization has shaped the classical but largely revisited hematopoietic model, in which the long-term HSC (LTHSC), at the top of the hierarchy, undergoes a lineage commitment through a series of discrete intermediate progenitor stages in a stepwise manner. This approach has help to categorize short-term self-renewal HSC (STHSC) and multipotent progenitor populations (MPP2, MPP3 and MPP4) ^3; 4; 5; 6^.

It is now evident that HSCs are not a homogeneous cell population and that each HSC does not contribute equivalently to all blood lineages: HSC heterogeneity was first suggested with single cell transplantation experiments showing that phenotypically identical HSC differs in self-renewal abilities and lineage differentiation potential ^7; 8; 9^; Next, single cell transcriptomic approaches combined with lineage tracing suggested an initiation of transcriptional lineage programs in HSCs, which bias their differentiation potential ^10; 11^ supporting an early HSC lineage segregation and a continuous differentiation model ^12^. Thus, it is now admitted that each individual HSC, although sharing the same marker combination, differs in terms of functional outputs and molecular signature ^13; 14; 15^.

This HSC heterogeneity has physiological consequences not only in terms of response to injury-induced infection and inflammation, which triggers emergency hematopoiesis and activates a subtype of phenotypic HSCs ^16; 17^, but it also intervenes upon physiological aging. Hematopoietic aging is associated with a reduced production of red blood cells and lymphocytes concomitant to an increase of myeloid and megakaryocytic cells, that promote immunosenescence and myeloid malignancies ^18; 19^. Evidence indicates that these alterations of the hematopoietic system come from an age-related modification of the HSC compartment. Intrinsic changes such as accumulation of DNA damage, changes in the activity of epigenetic modulators and imbalance between repressive and activating histone marks in HSCs have emerged as contributing factors of hematopoiesis aging ^20; 21^. HSCs that are heterogeneous with respect to their self-renewal and differentiation capacities at birth pass through clonal selection over time due to environmental cues ^22^. This results in an increase in myeloid- and megakaryocytic-biased but multipotent HSCs within the phenotypic LTHSC compartment ^23; 24^. Thus, aging is not only reflecting an intrinsic uniform change in lineage output of the HSCs but is rather due to a shift in the relative proportion of HSCs with different characteristics ^25^.

Previous studies on age-related transcriptomic changes of HSCs at the single cell resolution have revealed an expansion of platelet-primed HSCs ^26^ and a gain of a self-renewal expression program ^27^ with aging. However, the resolution of the analyses in particular regarding the proportion of the different HSC population and their variation upon aging were limited due to the small number of analysed cells and sorting strategies. Here, we took advantage of the 10× Genomics approaches and the development of new bioinformatic methods and tools to increase the resolution and revisit the transcriptional heterogeneity and change upon aging of the HSC compartment. By analysing 15,000 single murine hematopoietic stem and progenitor cells (HSPC) transcriptomes we could detect rare HSC subpopulations that accumulate upon aging. We could also highlight transcriptional program changes linked to cell cycle activity during aging that participate to the HSC age-related alterations.

## RESULTS

### Stratification of HSPCs using single-cell transcriptome analysis highlighted 15 different clusters

To characterize HSC populations by single cell RNAseq (scRNA-seq), we purified immature hematopoietic cells by FACS from BM pools of young (2-3 months) and old (17-18 months) mice. To catch very early events in the HSC differentiation process and isolate HSPCs, including LTSHCs, STHSCs, MPP2 and MPP3, we applied the widely used Lin^−^, Sca1^+^, cKit^+^ (LSK) marker strategy with the addition of the Flt3 marker to exclude the Flt3^+^ LSKs also referenced as MPP4 (Fig. 1a and Supplementary Fig. 1A). Four pools (2 pools of young and 2 of old) of thousands HSPCs were subjected to 10x Genomics Chromium capture platform and a total of 15000 single HSPC transcriptomes were sequenced (young pools, with 5189 and 2244 cells and old pools with 3328 and 4154 cells after quality control; Supplementary table 1). As we made the assumption that aging would not dramatically modify HSC identity, we first analysed young and old HSPCs together and performed batch and cell cycle correction using the Seurat workflow ^28^. Unsupervised clustering grouped the transcriptomes into 15 distinct clusters, which were visualized by Uniform Manifold Approximation and Projection (UMAP) ^29^ (Fig. 1b). We identified cluster markers using differential expressed gene (DEG) analysis (Supplementary Table 2) and cluster characteristics were deduced (Fig. 1c) from Gene Ontology term enrichment analysis (Supplementary Table 3) and confirmed with gene signatures related to hematopoiesis (Supplementary Table 4a). Six clusters were classified as lineage-primed clusters as they were clearly enriched for HSPCs with megakaryocyte (pMk), erythroid (pEr), neutrophil (pNeu), mastocyte (pMast), B lymphocyte (pB) or T lymphocyte (pT) commitment gene markers (Fig. 1b-d; Supplementary Tables 2, 3 and 4a). Nine clusters were considered as non-primed due to their lack in expression of lineage restricted-genes. They accounted for a large majority of the analysed cells (90%) (Fig.1b-d; Supplementary Tables 2 and 4b).

**Fig. 1.**
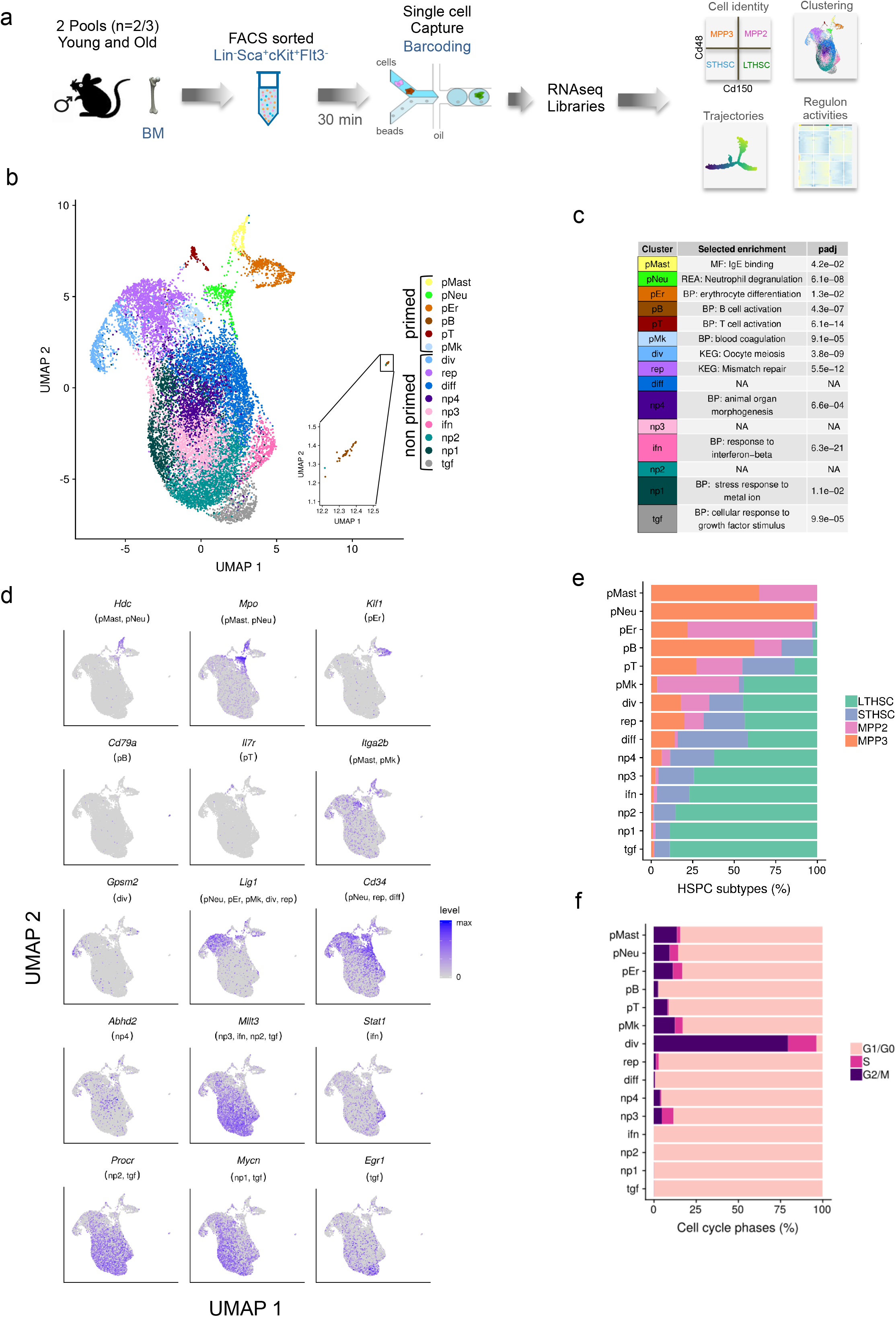
Unsupervised clustering of young and old HSPCs revealed 15 clusters gathering mainly immature and to a lesser extend lineage-primed HSPCs. **a** Overview of the scRNA-seq sample preparation and analysis. Cells were isolated from bone marrow (BM) of young and old mice and pooled to obtain 2 pools for each age. Pools correspond to 2 or 3 BM (n=2/3). BM cells were FACS sorted to purify Lin^−^, Sca-1^+^, c-Kit^+^ (LSK) Flt3^−^ cells that defined the HSPCs. The four pools of HSPCs were processed using droplet-based single cell sequencing (10X Genomics) and multiple analyses were performed using bioinformatics tools to characterize aging effects. **b** UMAP plot of young and old HSPCs (15000 cells) analysed using Seurat. Colours marked the 15 distinct clusters identified by gene enrichment analysis. Each dot represents a cell. **c** Selected enrichment for each cluster and corresponding p-values adjusted for multiple testing (padj). NA indicates non-relevant enrichment. **d** UMAP plots coloured by expression of selected cluster markers. Cluster names are indicated in parenthesis. **e** Percentage of LTHSCs, STHSCs, MPP2 and MPP3 within the HSPC population, identified by transfer learning in each of the 15 clusters. **f** Percentage of computationally assigned cell cycle (G1/G0, S and G2/M) phases in each of the 15 clusters.

To further characterize the clusters, we looked at the distribution of the 4 phenotypically distinct HSPCs, LTHSCs, STHSCs, MPP2 and MPP3, within these clusters. We first identified these four HSPC subtypes in our dataset by transfer learning using previously published scRNA-seq data coupled with FACS-labelled HSPCs ^12^ (Supplementary Fig. 1B) and then analysed their proportion per cluster. This showed that globally lineage-primed clusters were enriched with MPP2 and MPP3, suggesting their “more differentiated” state in comparison to the remaining clusters (Fig. 1e and Supplementary Table 4c). Interestingly, the neutrophil-biased cluster (pNeu) was almost exclusively enriched with MPP3 (98%), while pMast and pEr were enriched with both MPP2 and MPP3 (Fig. 1e and Supplementary Table 4c). Noticeably, the pMK cluster was composed of almost 50% of LTHSCs, supporting previous work suggesting that platelet-biased stem cells reside at the apex of the HSC hierarchy ^30^. Analysis of computationally assigned cell cycle phases in each of the 15 clusters confirmed that the “div” cluster was composed of dividing cells and indicated that globally the lineage-primed clusters contained a reduced proportion of cells in G1/G0 in comparison to the other clusters (Fig.1f and Supplementary Table 4d). This goes in line with their enrichment in MPP sub-populations and reflects their activated states.

Among the non-primed clusters, the four named np1, np2, np3 and np4 were overlapping and positioned at the centre of the UMAP (Fig. 1b). They could not be distinguished with specific gene expression signature (Fig. 1c, d; Supplementary Table 2 and Fig. 2) and were characterized by a high percentage of cells indexed as LTHSCs (Fig. 1e and Supplementary Table 4c). By contrast, 2 clusters, also composed mainly of LTHSCs, harboured a very distinguishable signature for growth factor signalling (tgf) and interferon response (ifn) respectively (Fig. 1b-d and Supplementary Table 3), witnessing the existence of cells with signalling features at the top of the differentiation hierarchy. The remaining 3 clusters (diff, div and rep) were composed of less than 50% of LTHSCs (Fig. 1e and Supplementary Table 4c) suggesting their intermediate state in term of differentiation. The cluster named diff had very few distinguishable markers but was enriched with Cd34 expressing cells (Fig. 1d and Supplementary Fig. 2). Interestingly, this cluster was the most enriched with STHSCs (Fig. 1e and Supplementary Table 4c), which have been characterized by the appearance of the Cd34 at their surface ^2^. The div cluster, characterized by enrichment for the Oocyte meiosis KEGG pathway (Fig. 1c and Supplementary Table 3) and genes involved in asymmetric division such as *Gpsm2* (Fig. 1d and Supplementary Fig. 2) was particularly different from the other clusters by its enrichment in G2/M cells (Fig. 1f and Supplementary Table 4d). The rep cluster was characterized by replication and reparation gene signatures and presented a specific high expression of *Lig1* (Fig. 1c, d and Supplementary Fig. 2 and Table 3).

**Fig. 2.**
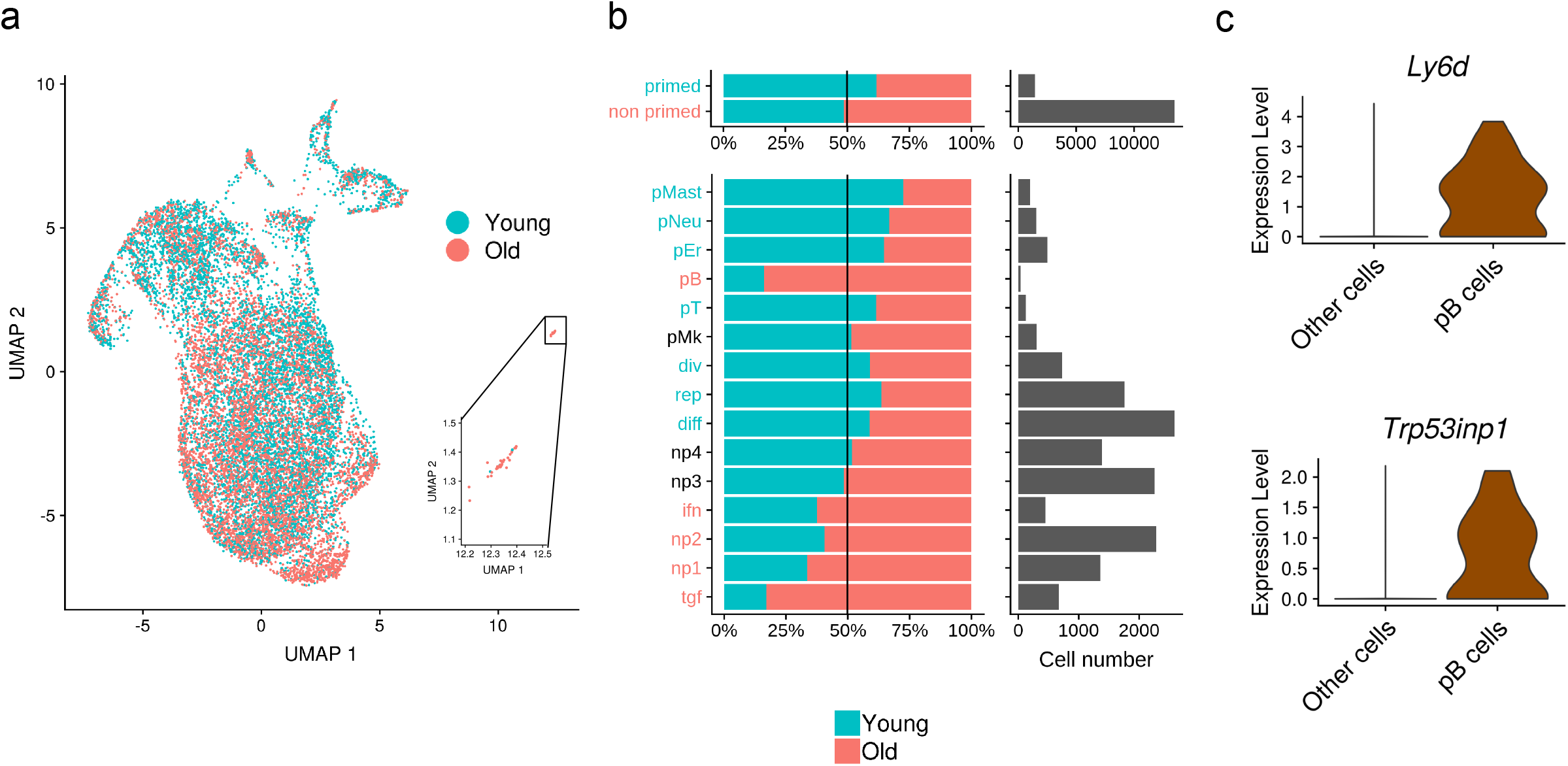
Aging affects more the immature than the lineage-primed HSPCs. **a** UMAP plot (same as in figure 1b) showing the young (blue) and the old (red) HSPCs. **b** Distribution of young (blue) and old (red) HSPCs in the clusters. On the left, percentage of young and old HSPCs in primed/non-primed clusters gathered and in each of the 15 clusters is presented. The Black vertical line indicates expected young and old cell proportions according to dataset size. Names of the clusters for which proportion of old or young cells was significantly higher than expected (hypergeometric test p-value < 0.05) are coloured in red for old cells and in blue for young cells. On the right, barplots represent the number of cells composing each ensemble: primed/non primed clusters gathered and individual clusters. **c** Violin plots showing *Ly6d* and *Trp53inp1* expression significantly up regulated in the pB cells cluster in comparison to the other cells (p-value < 0.05 & log fold change > 0.25).

As a whole, these results highlight the interest of gene expression signature to identify heterogeneity in the HSC population. They support the presence of differentiation-biased cells in the immature hematopoietic compartment and demonstrate that transcriptional programs can subdivide HSPCs in different clusters besides their classical differentiation state defined by cell surface markers.

### Aging affects HSPC clusters differently

Aged HSCs have been characterized by changes at the transcriptome level that could be the result of a shift of HSPC populations with different transcriptomic programs and/or of intrinsic gene expression changes ^20^. Yet, the relationship between these two aged-related changes is still poorly described. Here, we used our 15000 single young or aged HSPC transcriptomes to analyse age-dependent population in relation to gene expression modifications.

To assess the aging effect at the level of HSC populations, we first confirmed by FACS analyses and by transcriptomic based cell population predictions, the well-described accumulation of LTHSCs that occurs at the expense of the STHSCs and the MPP3 upon aging (Supplementary Fig. 1C, D). Analysis of young *versus* old cells in the UMAP plot showed that old cells were significantly more distributed in the non-primed clusters while lineage-primed clusters were enriched with young HSPCs (Fig. 2a, b). Indeed, the primed T-cell (pT) and the myeloid primed pMast, pNeu and pEr clusters were predominantly composed of young cells (Fig. 2b and Supplementary Table 4e). An exception was observed for the primed B-cell (pB) cluster; although representing very few cells, this cluster was comprised mainly of old ones (Fig. 2b and Supplementary Table 4b and e). Interestingly, these old B-biased cells were characterized, in addition to the expression of early B-cell markers such as *Ly6d* and *Cd79a* (Fig. 1d, 2c and Supplementary Tables 2), by *Trp53inp1* expression, for which we recently showed its involvement in the blockage of early B-cell developmental step ^31^. Thus, this cluster may represent an aged B-cell population that cannot complete its maturation. In addition to the pB cluster, four non-primed clusters, the np1, np2, ifn and tgf clusters were significantly enriched with old cells (Fig. 2b and Supplementary Table 4e), suggesting that aging favoured the amplification of specific featured LTHSCs. This result highlights an amplification of LTHSCs deregulated in their response to different stimuli such as inf and tgf signalling and, therefore, supports the previous observation of an increase of HSPCs with self-renewal and quiescence potential in older BM ^25^. Noticeably, amplification or reduction of a given cluster was not observed in the same way in our two batches of experiments. Indeed, we observed that the age induced decrease of pT cluster and increase of tgf cluster were mainly driven by one batch, specific for each of them (Supplementary Table 4e) witnessing a heterogeneity of aging inter mouse groups.

From these results, we conclude that, globally, aged haematopoiesis is stemming from HSPCs that are not lineage primed and that HSPCs/individuals are not affected equally by aging. These observations are of particular interest for the myeloid bias of aging haematopoiesis, which, according to our analyses, would not come from the amplification of cells with myeloid lineage priming (eg: pNeu or pMast) but could come from the amplification of non-primed HSCs.

### Gene expression is more altered upon aging in non-primed clusters, with a loss of differentiation and a gain of hemostasis signatures

To reveal age-dependent changes in gene expression, we first compared the transcriptomes of young and old HSPCs. Differentially expressed gene (DEG) analysis highlighted a global HSC aging signature that was characterized by an up regulation of the stress gene *Nupr1*, the platelet-lineage markers *Vwf* and *Clu*, and markers of undifferentiated HSPCs such as *Procr* and *Slamf1*, as well as by a down regulation of genes that mark HSC differentiation, such as *Cd34* and *Cd48* (Supplementary Fig. 3 and Supplementary Table 5). These results are in line with the altered differentiation potential and platelet bias of old HSPCs ^26^.

**Fig. 3.**
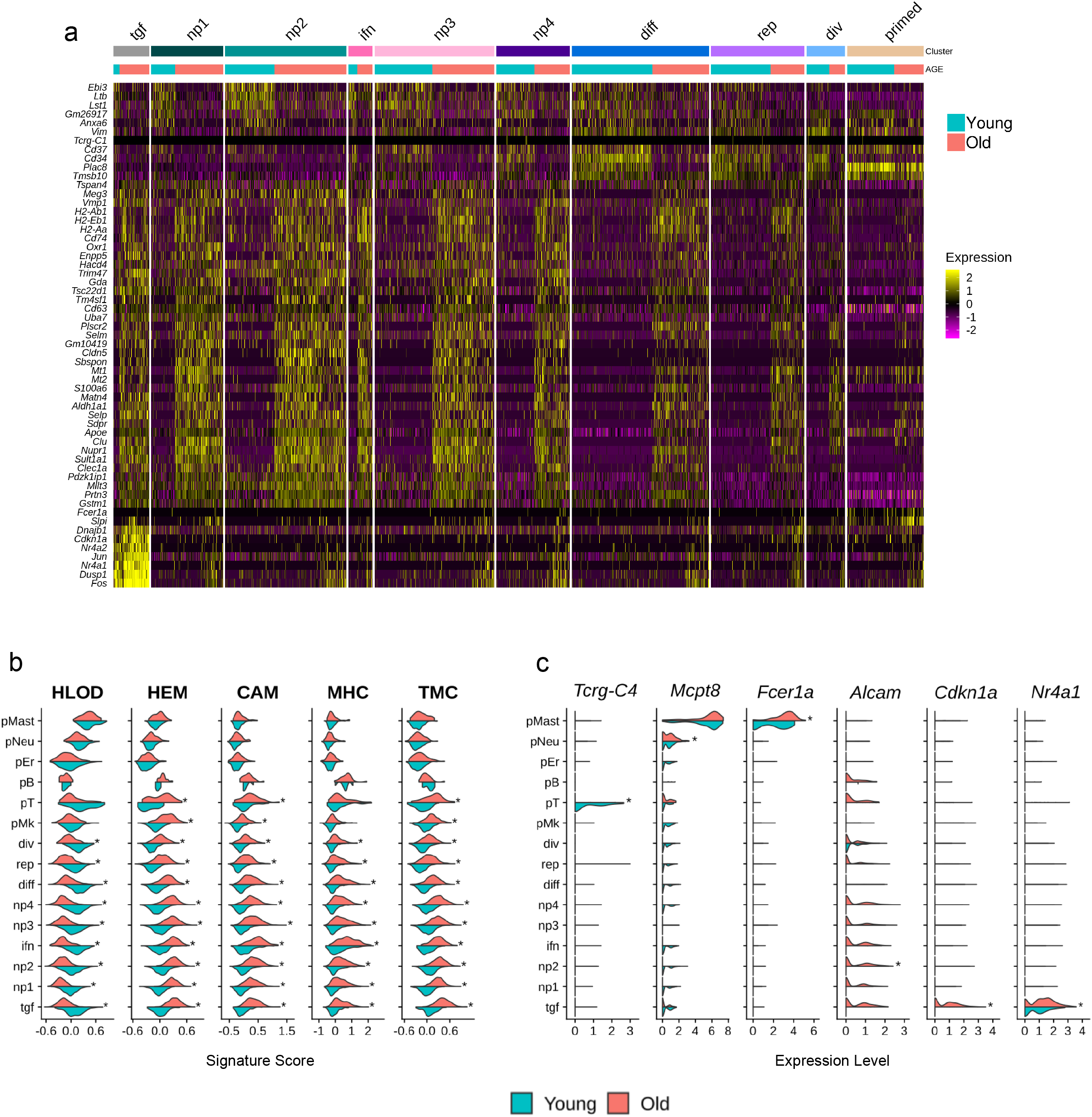
Gene expression is more impaired during aging in non-primed clusters with loss of differentiation and gain of hemostasis signatures**. a** Heatmap of the most significant differentially expressed genes (DEGs) during aging (p-value < 0.05 & log fold change > 0.5 in at least one cluster) in the different clusters revealed by Seurat analysis (Figure 1b). The lineage-primed clusters are gathered and labelled as primed. The upper coloured bars indicate cluster identity according to the colour code in figure 1b. The lower coloured bars indicate the proportion of young (blue) and old (red) cells in a given cluster. Gene expression are standardised across the entire dataset. **b** Combined violin plots showing signature scores (x-axis) in young (blue) and old (red) conditions per cluster. Signature scores represent the global expression of annotated genes for selected terms from enrichment analysis issued from DEGs during aging (p-value < 0.05 & log fold change > 0.25 in at least one cluster). Significant terms (enrichment gprofiler p-value < 0.05) are: Hematopoietic or Lymphoid Organ Development (HLOD) retrieved from GO:Biological Process, Hemostasis (HEM) retrieved from REACTOM pathways, Cell Adhesion Molecule (CAM) retrieved from KEGG pathways, MHC protein complex (MHC) retrieved from GO:Cellular Component, Transcriptional Miss-regulation in Cancer (TMC) retrieved from KEGG pathway. See supplementary Tables 7A & B for the lists of genes enriched in the terms. Stars show significant differences between the signature scores of young and old cells, per cluster (average score difference > 0.1 and p-value < 0.05). **c** Combined violin plot showing aging marker expression in young (blue) and old (red) conditions for the different clusters. Stars show significant differences of gene expression between young and old cells (average log fold change > 0.25 and p-value < 0.05).

In order to assess the heterogeneity of transcriptome changes upon aging according to HSC clusters, we analysed changes in gene expression of each cluster separately (Supplementary Table 6). Heatmap of the most differentially expressed genes (DEGs) (log fold change > 0.5) upon aging analysed per clusters showed that the non-primed clusters exhibited more differences in their transcriptome than the primed ones and that these differences were towards an increase of gene expression rather than a decrease, suggesting an increased cell-to-cell transcriptional variability upon aging (Fig. 3a and Supplementary Fig. 4). For these non-primed clusters, except for the tgf cluster, the differential gene expression analysis per cluster followed the aging signature that was observed when analysing the totality of the cells (R^2^ > 0.8 Supplementary Fig. 5). Enrichment analysis of DEGs upon aging revealed a negative regulation of hematopoietic or lymphoid organ development (HLOD) marked by the down regulation of *Cd34*, *Plac8* and *Foxo3* (Supplementary Table 7A), together with a positive regulation of hemostasis with *Clu* and *Selp* increased expression, Cell Adhesions Molecule (CAM) genes such as *Alcam, Jam2,* Major Histocompatibility Complex (MHC) H-2 genes and genes involved in transcriptional mis-regulation in cancer (TMC) (Supplementary Table 7B). TMC enrichment, in addition to TFs such as *Fli1* and *Pbx1*, relies on cell cycle kinase inhibitors *Cdkn1a* and *Cdkn2c* and the stress response gene *Nupr1* suggesting a deregulation of the cell cycle phases upon aging. Globally, we found that aging feature-score differences were more pronounced in the non-primed clusters than in the lineage-primed ones (Fig. 3b Supplementary Table 7C). However, this analysis per cluster highlighted that among the lineage-primed clusters the pT and pMK clusters were still transcriptionally affected by aging, with an increase in HEM, TMC and CAM signatures (Fig. 3b and supplementary Table 7C). Looking at some genes individually, we were able to highlight some age-related changes affecting peculiar clusters. We observed a downregulation of the T-cell gene *Tcrg-C4* in the old pT cluster and an upregulation of the protease mast cell gene *Mcpt8* and myeloid integrin gene *Fcer1a* in old pMast and pNeu cluster respectively (Fig. 3c supplementary table 6). We observed an upregulation of *Alcam* required for HSC maintenance in np2 clusters (Fig. 3c and supplementary Table 6). Finally, we also observed a very specific transcriptome in old tgf cluster characterized by an increase of genes involved in HSC quiescence such as *Cdkn1a*, *Nr4a1* (Fig. 3c), which were clustered together in the heatmap of DEGs upon aging (Fig. 3a).

**Fig. 4.**
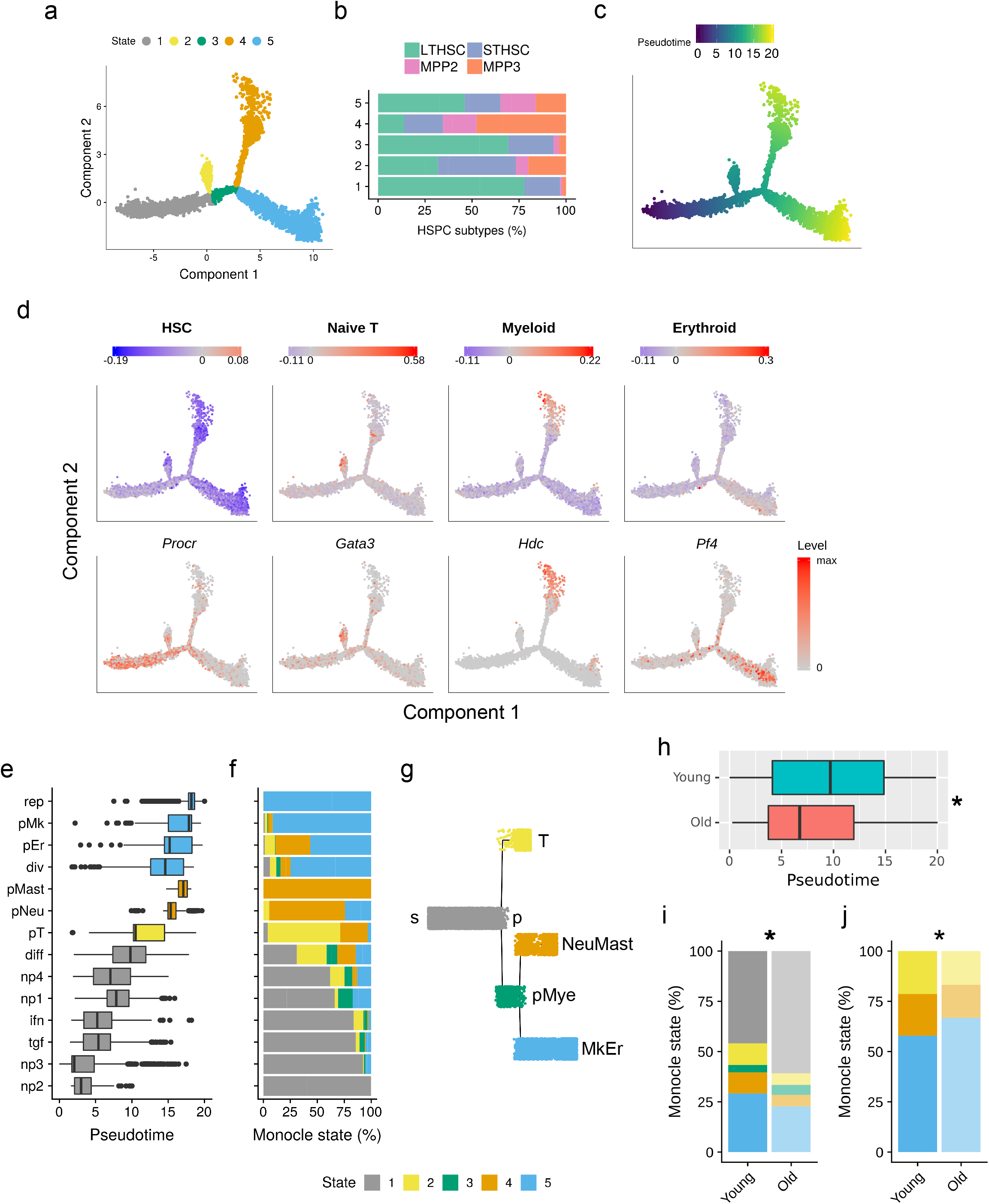
HSPC differentiation trajectory revealed a clear split between T-, MastNeu and MkEr primed cells. **a** Differentiation trajectory generated using Monocle 2 with all HSPCs excepted primed B cells that were excluded. Cells are coloured according to five states (1 grey, 2 yellow, 3 green, 4 orange and 5 blue), which partition the trajectory. **b** Barplots representing the LTHSC, STHSC, MPP2 and MPP3 proportions in the five states. **c** HSPC differentiation trajectory coloured according to HSPC pseudotime values and representing their differentiation progression. **d** HSPC differentiation trajectories coloured according to HSPC scores for hematopoietic lineage signatures retrieved from the literature (upper panel) and according to the expression level of HSPC differentiation markers (lower panel). For signatures, positive (red) or negative (blue) scores indicate whether the expression of the tested genes is more or less important than the reference signature. Signatures identified are HSC, naïve T, Myeloid and Erythroid. HSPC differentiation markers shown are: *Procr* for LTHSCs, *Gata3* for T-cell primed HSPCs, *Hdc* for myeloid-primed HSPCs and *Pf4* for erythrocyte-megakaryocyte-primed HSPCs**. e** Repartition of the Seurat clusters along the pseudotime. Box plots of pseudotime values are coloured according to the most represented state. **f** Repartition (in percentage) of the different states (1 to 5) of the trajectory for each Seurat cluster. **g** Tree representation of HSC differentiation trajectory, edges representing the states (state 1 in grey, 2 yellow, 3 green, 4 orange and 5 blue), and nodes standing for pseudotime points: the starting point (s), the first bifurcation point (p), the primed Myeloid bifurcation point (pMye); and the three fates T-lymphocyte (T), Neutrophils/Mastocytes (NeuMast) and Megakaryocyte/Erythroid (MkEr). **h** Boxplot of pseudotime value for young and old cells. * indicates a significant difference between young and old pseudotime value distribution (median difference > 2.9, p-value < 10^−16^ Wilcoxon rank sum test) **i - j** Percentage of Monocle states in young and old conditions, when considering all states (i) or only states 2, 4 and 5 (j). * indicates a significant dependence between state and age repartitions (p-value < 10^−10^ Pearson’s Chi-squared test).

**Fig. 5.**
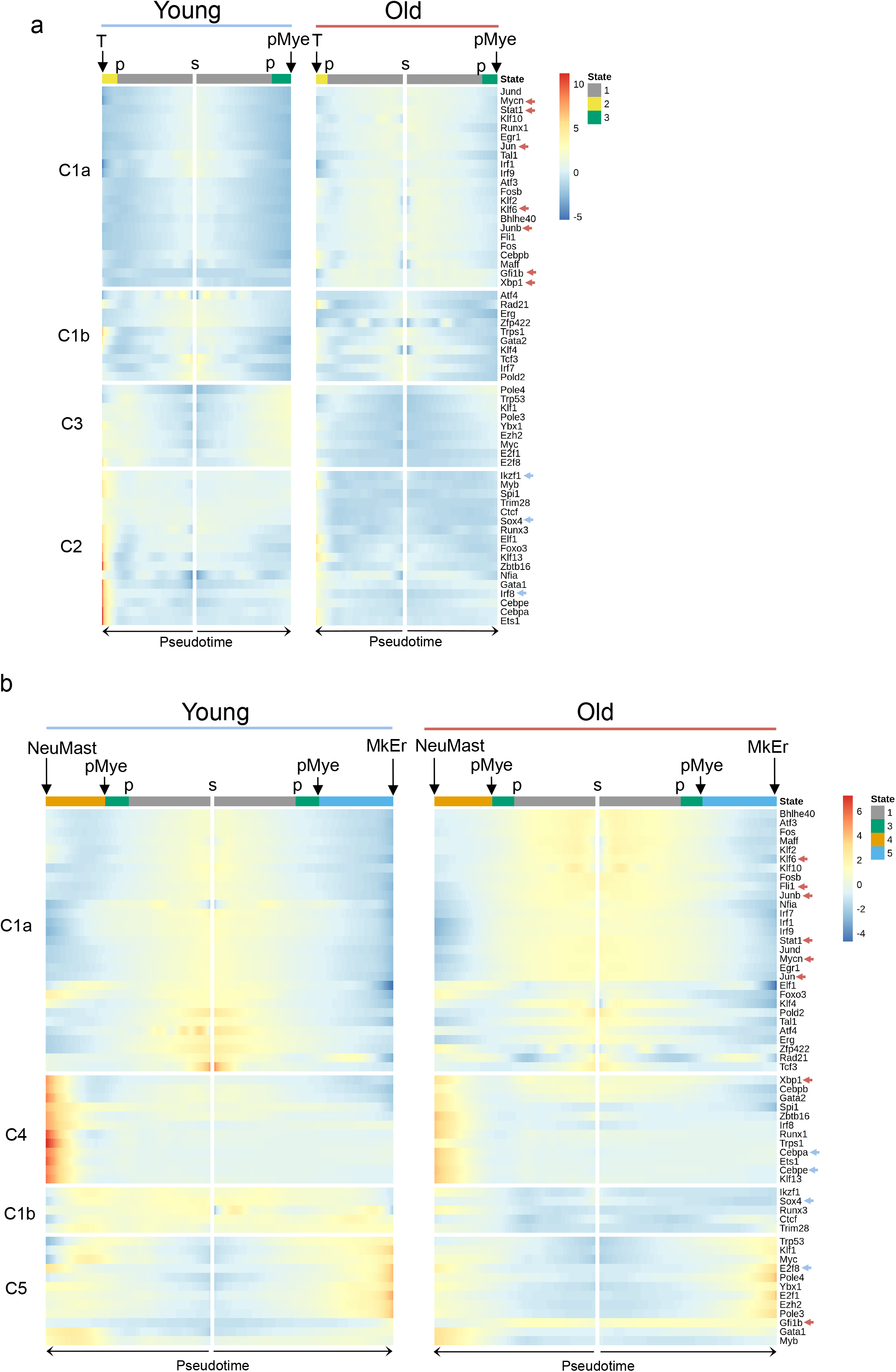
HSPC differentiation trajectory associates with transcriptional programs that are altered upon aging. **a**-**b** Heatmaps showing standardised regulon activity scores, recovered with the AUCell procedure of Scenic, for young (left panel) and old (right panel) HSPCs across Monocle states. Cells (columns) were ordered according to their pseudotime, and colour bars at the top of the heatmaps indicate the state at which cell belongs (1 grey, 2 yellow, 3 green, 4 orange and 5 blue). Regulons (rows) were hierarchically clustered, based on their activity score in young HSPCs. In **a,** 4 clusters of regulons are highlighted when analysing regulon activity along pseudotime trajectories from s to T fate and from s to pMye bifurcation point (i.e. across Monocle states 1, 2 and 1, 3). In **b,** regulon activity along pseudotime trajectories from s to NeuMast and from s to MkEr fates (i.e. across states 1, 3, 4 and 1, 3, 5) is analysed and 4 other clusters of regulons are highlighted. Arrows mark regulons for which a significant difference of activity with aging (average AUCell score difference between young and old cells > 0.002 and p-value < 0.05) were found in at least one of the considered states (i.e states 1, 2 and 3 in **a** and 1, 3, 4 and 5 in **b**). The colour indicates if regulon activity is increased (red) or decreased (blue) in old compared to young cells.

Altogether, our results pointed peculiar age-related changes mostly affecting the transcriptome of HSPCs from non-primed clusters and characterized by a loss of differentiation genes that could account for the functional changes of the aged hematopoietic compartment.

### Differentiation trajectory shows a HSPC progression toward T, Mast/Neu and Mk/Er fates that is altered with age

It has been recently suggested that HSCs undergo a continuous differentiation process rather than a stepwise process ^13^. In order to better capture and understand the progression of this differentiation process during aging, we constructed pseudotime trajectories by ordering HSPCs based on the similarities between their expression profiles with Monocle ^32^. We first generated the trajectories of young and old HSPCs separately. The two resulting trajectories showed a very similar shape, with the exception of a group of cells standing apart from the old trajectory, and made exclusively of pB cells (Supplementary Fig. 6A). Because these cells were detected only in old HSPCs and were clearly distant from the rest of the cells in the UMAP (Fig 1b), we excluded them for cell pooling and ordering for both ages. Thus, we analysed the differentiation trajectory inferred from young and old cells pooled together, without pB cells. The resulting trajectory was partitioned into 5 segments, called Monocle states labelled states 1, 2, 3, 4 and 5 (Fig. 4a). The departure of the trajectory was identified at the extremity of the state 1, as this state possessed the highest percentage of LTHSCs (Fig. 4b, c). States 2, 4 and 5 were enriched with MPPs suggesting their progression towards differentiated states (Fig. 4b, c). The 5 states of the trajectory were characterized with gene expression based on previously published signatures related to HSPCs and hematopoiesis (referenced in Supplementary Table 8A) and on our state marker analysis (Supplementary Table 8B). This characterisation revealed that HSPCs in state 1 expressed a HSC signature at a higher level than in the other states with especially cells expressing the dormant HSC marker (*Procr*); state 2 cells (after the first bifurcation) were characterized with Naive T-cell signature and were expressing *Gata3*, suggesting a primed-T differentiation state (Fig. 4d); state 4 cells were characterized by a myeloid signature ^33^ and high expression of *Hdc*, previously reported as a marker of myeloid biased HSPCs ^34^, while cells in state 5 presented an erythrocyte signature ^33^ and expressed *Pf4* a megakaryocytic marker (Fig. 4d). To better characterize state 2 and because this state shared 72% of its markers with state 4 (Supplementary Table 8B), we looked at DEGs between the two states, which highlighted an up regulation of genes involved in T cell differentiation such as *Ctla2a, Zfp36l2* and *Gata3* in state 2 (Supplementary Table 8C), confirming the primed-T identity of state 2 cells.

**Fig. 6.**
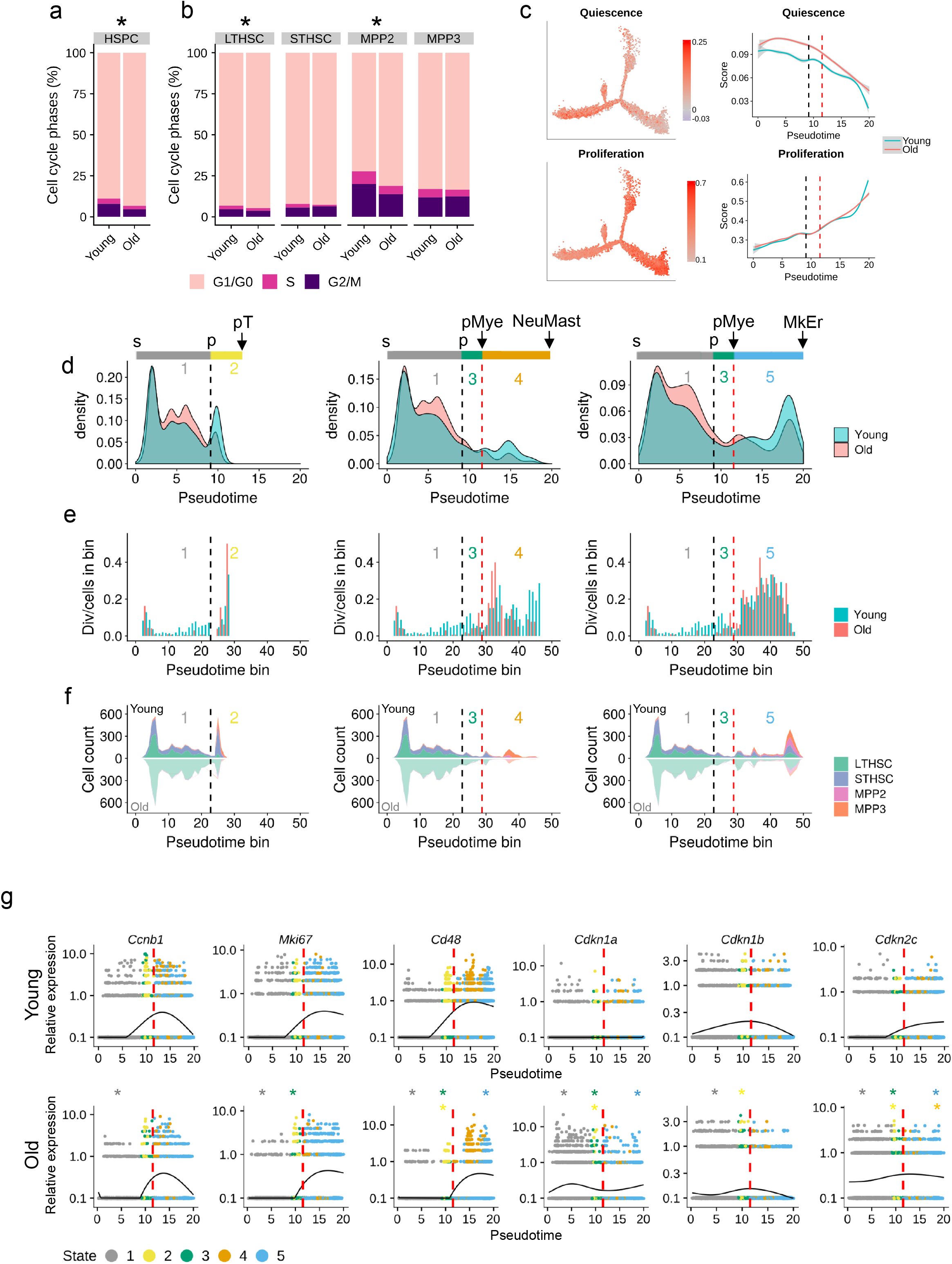
Cell cycle analysis along pseudotime highlights a delay in differentiation associated with cell cycle arrest in aged condition. **a** Repartition (in percentage) of the cell cycle phases (estimated with cyclone) in young and old HSPCs. **b** Repartition (in percentage) of the cell cycle phases (estimated with cyclone) in LTHSCs, STHSCs, MPP2 and MPP3 in young and old conditions. For a and b **\*** indicates a significant dependence between cell cycle phase and age repartitions (p-value < 0.05 Pearson’s Chi-squared test). **c** Left panel, differentiation trajectory of HSPCs coloured in accordance to their score for previously published quiescence and proliferation signatures; Right panel, comparison of the scores for the quiescence and proliferation signatures between young and old HSPCs in pseudotime. **d** Density plot of young (blue) and old (red) cells along pseudotime for the T (left), NeuMast (middle) and MkEr (right) fates. Black and red dashed lines mark respectively p and pMye bifurcation points. **e** Division rate along pseudotime for young (blue) and old (red) HSPCs for the T (left), NeuMast (middle) and MkEr (right) fates. On x-axis, pseudotime was cut into 50 bins and a division rate is calculated for each bin, by dividing the number of young (*resp*. old) cells assigned to G2M phase by the total number of young (*resp*. old) cells of the bin. Black and red stretched lines mark p and pMye pseudotime bifurcation point respectively. **f** Stacked plot of predicted cell types along pseudotime cut into 50 bins for young (upper part of the plots) and old (lower part of the plots), for the T (left), NeuMast (middle) and MkEr (right) fates. Black and red stretched lines mark p and pMye bifurcation point pseudotime respectively. **g** Relative expression of some genes as a function of pseudotime for young (upper panel) and old (lower panel) HSPCs. Points represent cells, which are coloured according to their belonging to the 5 different states (1 grey, 2 yellow, 3 green, 4 orange and 5 blue). The y-axis is in log scale. * indicates significant differences in gene expression between young and old cells (p-value < 0) and star colour indicates the state where the difference is found.

Analysis of Seurat cluster position and spreading on the trajectory (Fig. 4e; Supplementary Fig. 7) strengthened the pseudotime differentiation relevance with lineage-primed clusters located at the two extremities of the trajectory and suggested a differentiation specificity of the states (Fig. 4e). Study of the five state proportions across the clusters, revealed a first bifurcation separating pT cells (state 2), from cells primed for myeloid lineages (state 3), and then a clear branching between Neu/Mast-primed (NeuMast) HSPCs (state 4) and Mk/Er-primed (MkEr) HSPCs (state 5) (Fig. 4f). The specificity of state 5 for megakaryocyte differentiation was supported by the high representation of the rep cluster (Fig. 4f), characterized by a reparation gene signature (Supplementary Table 3), which was previously associated with megakaryocyte fate ^35^. Separate pseudotime ordering of young and old HSPCs provided very similar segregation between the lineage-primed HSPCs, with one bifurcation from LTHSC (state 6) towards Neu/Mast-primed (NeuMast) HSPCs (state 7) and Mk/Er-primed (MkEr) HSPCs (state 8) (Supplementary Fig. 6A-E). However, the bifurcation towards the T lymphocyte fate was not retrieved probably because of the reduction of the pT cell number due to the sample splitting (Supplementary Fig. 6A). Hence, to synthetize our analyses, we proposed a tree-representation of the HSC differentiation trajectory (Fig. 4g) where nodes stand for pseudotime points, and edges for Monocles states. It contains 6 nodes: a root, the starting point (s); two internal nodes, the first bifurcation point (p) and primed Myeloid bifurcation point (pMye); and three leave nodes, the three fates T-lymphocyte (T), Neutrophils/Mastocytes (NeuMast) and Megakaryocyte/Erythroid (MkEr).

Next, we compared the differentiation progression of young and old HSPCs. Old HSPCs appear to be significantly delayed in the pseudotime (fig. 4h) while Seurat cluster spreading along the trajectory showed no clear differences of any cluster pseudotime position according to age (no median difference higher than 0.8 unit of pseudotime; Supplementary Fig. 8A). Looking at the proportion of the different Monocle states of the trajectory according to age revealed an increase in old HSPCs in states 1 and 3 in comparison to young ones (Fig. 4i). The old HSPCs accumulating in state 1 and 3 belong to the non-primed clusters np3, tgf, ifn, np4, diff and div (Supplementary 8B), confirming the accumulation of old HSPCs in non-primed clusters, localized earlier in the pseudotime than the lineage-primed ones (Supplementary Fig. 7). When focusing on cells belonging to states 2, 4 and 5, which reflect the 3 lineage-primed HSPC states, we observed that the proportion of state 5 (MkEr fate) was larger in old than young condition (Fig. 4j), although age was not affecting the percentage of the Monocle states from lineage-primed cluster cells (Supplementary Fig. 8B). This suggests that while less aged HSCs were detected in the three differentiation paths, cells with MkEr fate are more maintained upon aging than the ones toward NeuMast and T fates.

In conclusion, our trajectory analysis revealed a priming of HSPCs for T lineage that occurs early in the differentiation process and evidenced a clear split between the NeuMast and the MkEr HSC fate identifying an early lineage specification of HSCs (Fig. 4g). While the global shape of the trajectory and the lineage specification of the HSPCs are conserved upon aging, repartition of the old HSPCs along the differentiation trajectory is altered with a decrease in terminal states 2 and 4 conducing respectively to T and NeuMast fates, in favour to cells of the initial states 1 and 3.

### HSPC differentiation trajectory is associated with transcriptional programs that are altered upon aging

Cell fate decision and proper function of HSCs rely on tightly controlled transcriptional programs orchestrated by transcription factor (TF) activity ^36^. Since level of the expression of TFs is not sufficient to assess their activity, we measured changes in TF activity during differentiation and aging of HSPCs. For that, we took advantage of Single-Cell Regulatory Network Inference and Clustering (SCENIC) approach ^37^ that calculates the activity of a given TF (regulon score) based on target expression and cis-regulatory elements. We considered 154 TFs, selected from the literature or from our single cell expression data analysis (Seurat cluster markers), out of which, 58 were identified as active regulons in our HSPCs (Supplementary Table 9A). By looking at regulon activities of young HSPCs along the trajectory, we revealed a specific regulon signature for each state (Supplementary Table 9B). State 1 was characterized with activity of the stress sensors Atf3, the interferon signalling factors, Irf1, Irf7, Irf9 and the downstream targets of the Tgfbeta signalling, Stat1, Klf4, Egr1, Klf6, Junb, depicting a stemness state (regulon clusters C1a and C1b Fig. 5a and C1a Fig. 5b, young panel). Comparison of TF activities between state 2 and state 3 at the first bifurcation (p) emphasized the T fate of state 2 with the detection of high activity of the T-cell transcription factors Ikzf1, Sox4 (regulon cluster C2 Fig. 5a, young panel) while state 3 cells enter a more general differentiation program with a slight increase of regulon activities such as Myc (regulon cluster C3 Fig. 5a, young panel). As expected, aging reduced the activity of the two regulons in state 2 witnessing the reduced T-cell activity during aging. By contrast, Klf6, Junb, Jun and Stat1 activities of old HSCs were spread and increased in old states 1 and 3, (Fig. 5a, b Supplementary Table 9C), which was consistent with the stem cell activity of old states 1 and 3 containing mainly LTHSC (fig 4b).

By looking at the second bifurcation (pMye) between state 4 and state 5, we confirmed that state 4 was neutrophil- and mast-biased as it was indorsed with a high activity of C/ebpa-e, Runx1 and Irf8, involved in myeloid differentiation (regulon cluster C4 Fig. 5b, young panel). Noticeably, aging decreased the activity of regulons involved in myeloid fate such as Cebpa and -e in state 4 (Fig. 5b and Supplementary Table 9C). This result was consistent with the decrease of Neutrophils and Mastocyte primed-cell number with aging (observed in fig 2B) and strengthened our hypothesis that myeloid bias of aged haematopoiesis, would not come from this path of the trajectory. Cluster C5 of the heatmap shows that State 5 was characterized with a strong activity of Klf1, E2f8, Ybx1, Gfi1b and Ezh2, all of which are implicated in the erythroid/megakaryocyte development (regulon cluster C5 Fig. 5b, young panel). Interestingly, the activity of E2f8 was significantly reduced with aging in state 5 whereas Gfi1b activity was considerably increased in this old state. It should be noted that Gfi1b is the regulon that experienced the greatest increase in activity with aging, not only in state 5, but also in all states where the greatest increase observed was in state 1. As Gfi1b is master regulator of thrombopoiesis (reviewed in ^38^) and as we found some of its targets such as *Clu, Esam* and *Serpinb1a*, annotated for hemostasis (Supplementary Table 9A) upregulated with aging (Supplementary table 7B), we suggested that Gfi1b sustains the platelet bias of old HSPCs.

Thus, TF activity analyses over the pseudotime corroborated the trajectory features and clearly identified a separation in TF activity that explains the T-lineage priming (Fig. 5a) and the two distinct myeloid fates, NeuMast and MkEr (Fig. 5b). It also indicated that aging is associated with marked changes in TF expression and activity with a gain of TFs involved in stemness and platelet activity and a loss of lineage-specific factors that drive lineage commitment and terminal differentiation.

### Cell cycle analysis along pseudotime highlights a delay in differentiation associated with cell cycle arrest in aged condition

As one of the hallmarks of HSC aging is a reduction of cycling HSCs ^39^ we analysed the cell cycle phases according to BM age. We showed an increase of non-cycling HSPCs (G1/G0) at the expense of the S and G2/M phases in old BM in comparison to young one (Fig. 6a). Analysis of LTHSC, STHSC, MPP2 and MPP3 population separately showed that age did not typically affect the proportion of cycle phases within each subtype, with the exception of a slight but significant change in LTHSCs and MPP2 (Fig. 6b). This suggests that the increase of the G1/G0 phase proportion observed upon aging is mainly due to the accumulation of quiescent LTHSCs that are known to be arrested in G1/G0 phase ^40^, and to a lesser extend to LTHSC and MPP2 intrinsic cell cycle changes induced by aging.

Previous studies have pointed a clear link between HSC cell cycle and commitment ^41^. Positioning quiescent versus proliferative cells along the trajectories showed that quiescent cells were at the departure of the trajectory while proliferating cells were towards the differentiated states (Fig. 6c, left panel). Comparison of the quiescence and proliferation signatures between young and old HSPCs showed a quiescence gain in the old condition in the first part of the trajectory (states 1, 2 and 3) while the proliferation signature remained unchanged (Fig. 6c, right panel and supplementary Table 10A).

Next, we addressed the question of the cell cycle and its influence on HSPC aging. We first analysed the distribution of young and old HSPCs along the trajectory, analysing T, NeuMast and MkEr fates separately (Fig. 6d). Doing so, we confirmed the accumulation of old HSPCs in state 1 before the first bifurcation point p and the decrease of old cells in the differentiated states 2, 4 and 5 (Fig. 6d). To associate cell-cycle status and cell accumulation, we performed a high-resolution analysis of cell cycle progression along the trajectory by plotting the ratio of dividing cells on pseudotime bins for young and old cells in T, NeuMast and MkEr fates separately (Fig. 6e). This highlighted a dramatic loss of dividing cells in old condition in state 1 with the exception of cells located at the very beginning of the trajectory (Fig. 6e). We hypothesised that these dividing cells (that are LTHSCs and belong to np3 cluster) represent cell-cycle activity of self-renewing LTHSCs. Interestingly, we found no difference in cell cycle phase proportion between these young and old LTHSCs (p-value > 0.3 Pearson’s Chi-squared test Supplementary Fig. 10), suggesting a conservation of self-renewal potential in old HSCs. By opposition, the absence of cell cycle activity of old HSPCs later in state 1, which may represent cell cycle activity linked to differentiation, underlines a default in cell division of old HSPCs associated to differentiation (Fig. 6e). Division rate of old HSPCs became positive after the first bifurcation and was similar to what we observed in young HSPCs (Fig. 6e), with the exception of a decrease in old cycling cells in state 4 (toward NeuMast fate) suggesting a default of cell cycle in old Neu-primed HSPCs. Visualization of the distribution of the different HSPC subsets confirmed the accumulation of old LTHSCs at the expense of the STHSCs and revealed a dramatic loss of NeuMast-primed cells upon aging (Fig. 6f).

We took advantage of our analysis of DEGs with aging per monocle state (supplementary table 10B) to identify DEGs involved in proliferation and cell cycle as well as in differentiation and analysed their expression profile in young and old cells along the trajectory. For the two proliferation-division genes, *Ccnb1* and *Mki67,* we observed a pronounced increase of expression in the young HSPCs that was occurring in state 1 concomitant to the increase of the marker of differentiation *Cd48* (Fig. 6g). In the old cells, increase in the expression of these three genes was also detected but was delayed until the branching point pMye suggesting a delay in the commitment of old HSPCs. To grasp molecular mechanism(s) that could be involved in this delay, we compared cell cycle inhibitor expression across young and old HSPC trajectory. *Cdkn1a* and *Cdkn2c* were upregulated along the old trajectory (except in state 4 for Cdkn2c) especially in the first part of the trajectory (state 1, 2 and 3) in comparison to young one. By contrast, Cdkn1b was downregulated in state 1 and 2 of the old trajectory (Fig. 6g and Supplementary table 10B). The change in expression with aging of the three cell cycle inhibitors known to control the fate of HSCs indicates deregulation of cell cycle progression in aged HSCs. It is interesting to note that *Cdkn1a* was found to be a target of Stat1, Jun and Junb which are themselves targets of the Klf6 regulon (Supplementary Table 9A), four regulons whose activities increased with aging in the same range of pseudotime as changes in the level of *Cdkn1a* expression (Fig. 6g and 5b).

Together, these results suggest that aged HSCs have a default in cell cycle, concomitant to a delay in their differentiation program, which occurs before the lineage priming of the HSPCs.

## DISCUSSION

HSCs represent a heterogeneous cell population in terms of their ability to self-renew and differentiate. In this study we questioned the effect of aging on HSC populations and properties. As scRNAseq now provides a powerful method for defining cell subtypes as well as a detailed description of the functional properties specific to these subtypes, we profiled 15,000 mouse HSPCs from young and old mouse BM by scRNAseq. This high resolution profiling of young and old HSPCs was the base to generate a reference map of mouse HSPCs and understand how this map is affected during aging.

At first, the large number of cells analysed provided us new insights of HSPC heterogeneity, through the identification of 15 distinct HSPC clusters that we divided in two categories, the non-primed clusters by opposition to the lineage-primed clusters composed of low-abundant HSPCs with restricted lineage potential that was previously described ^12^. Indeed, our analyses identified distinct lineage-primed HSPCs; HSPCs not only with Mk–restricted signature, which were widely reported within the HSPC compartment ^9; 30; 12^ but also HSPCs with mastocytes, neutrophils, erythrocytes, lymphoid B and lymphoid T-restricted lineage signatures. The wide range of lineage potentials of HSPCs detected in this study favours an early HSPC uni-lineage segregation ^42; 43; 12^ and argues against the previous notion of a unique HSC with multi-lineage potential ^44^. By using pseudotemporal reconstruction of differentiation trajectories, we further investigated the early pre-determined HSPC potential by highlighting bifurcations in the trajectory that reflect clear separations in the fate of specific lineage-primed HSPCs. Our clustering analysis in addition to transcriptional activities detected along the trajectories clearly characterised two bifurcation points, revealing three distinct HSPC fates towards T lymphocyte, Neu/Mast or Mk/Er lineages. In addition, our analysis showed that lineage priming of HSPCs is not delineate by a specific HSPC subset such as LTHSC, STHSC, MPP2 and MPP3 in any instance. Although we showed that Neu priming is clearly stemming from MPP3 and Er priming from MPP2, lineage priming could also arise from a combination of HSPC subsets. For example, the megakaryocyte potential was stemming from LTHSC and MPP2 subsets, in line with previous studies ^12; 26^. In addition, we detected a T potential in the four subsets of HSPCs and a B potential in MPP2 and MPP3, suggesting a lymphoid-priming occurring earlier in the BM and not restricted to the more engaged Flt3 positive MPP4 as previously reported ^45^. The fact that we detected lineage primed cells in the very early subset of HSPCs goes in line with a previous study showing the existence of four distinct and closely related stages of self-renewing LTHSCs in adult BM that stably adopt lineage-restricted fates (platelet, B and T lymphoid, erythroid and myeloid lineages) despite remaining multipotent ^46^.

If the accumulation of very immature HSCs in the BM of aged individuals is now an accepted criterion of hematopoietic aging, we still do not fully understand what are the characteristics of these aged HSCs and what causes them to accumulate. In terms of characteristics, it has been reported that accumulating old HSCs are LTHSCs with platelet-restricted lineage output^26^. However, this feature was challenged by a study showing that aging was characterized by the existence of latent-HSCs, a subpopulation of aged HSCs that displayed a five blood-lineage (Platelets, Erythrocytes, neutrophils-monocytes, T-, and B-lymphocytes) HSC phenotype following transplantation into secondary recipients ^24^. By looking at the transcriptomic changes at the single cell scale, we confirmed the global increase of the LTHSC fraction within the HSPCs. However, by analysing our old HSPCs by clusters or individually we could demonstrate that HSPCs are not affected uniformly by aging and grasp some interesting aging feature. At first, we showed that the proportion of old HSPCs in pMast, pNeu pEr and pT primed clusters was decreased while increased in ifn, tgf, np1 and np2 clusters. In addition young and old cells were found in the expected ratio in the pMk cluster. This clearly indicates that the platelet and myeloid bias observed upon aging is not due to an amplification of the pool of lineage restricted cells but stem from other subsets of HSPCs. Secondly, we highlighted some specific amplification of LTHSCs such as LTHSCs with miss-regulated interferon signalling (ifn cluster). As the increase in interferon response with aging in a number of different tissues has been observed ^47^ and is consistent with the concept of inflammaging ^48^, this amplification could afford for the myeloid bias observed in aging. Another interesting HSC group that we detected amplified during aging was the cluster of LTHSCs harbouring a Tgf signature that may correspond to the accumulation of the HSC subtypes with differential responses to TGF that was previously identified ^49^. These two types of old HSCs need to be further analysed but considering their characteristics it is tempting to hypothesize that their proportion was increased under stress selection pressures to compensate for the loss of mature cell production that occurs upon aging. They might witness the emergence of competitive clones that amplify during aging and fit quite well with the clonal haematopoiesis model. In another perspective, the apparition of the pB-primed cluster that we observed quasi exclusively in the old BM might also represent clonal evolution. Since this cluster was characterized with the expression of *Trp53inp1*, a gene limiting lymphoid differentiation upon aging, it could correspond to an accumulation of old HSPCs altered in their lymphoid differentiation ^31^ but resulting for a pressure of immune deficiency.

Pseudotime trajectory analysis led us to address the question concerning the differentiation state of old LTHSCs, which were thought to accumulate in a more undifferentiated state compared to young LTHSCs ^27^. First, the outcome of our analyses is in favour of no difference in term of differentiation state between young and old LTHSCs as when plotted together along the trajectory the most immature old cells were not positioned at an anterior pseudotime compared to the young ones. Second, our results support that old LTHSCs are delayed in their differentiation journey in comparison to young ones and that this delay occurs pretty early in the pseudotime, before the first bifurcation point that splits T fate from myeloid fate. This was clearly emphasised by our regulon activity analysis of transcription factors such as Myc, Trp53 or Spi1 that were previously described involved in multipotency and commitment of HSCs ^50^ and for which we could observe a delay in their activity along the differentiation trajectory.

Thus, the old HSCs are not more undifferentiated that the young ones but seem to have intrinsic defaults that would delay their commitment. This finding is interesting when putting in perspective what causes the accumulation of LTHSCs. Increase of LTHSCs with aging could originate from an increase in the self-renewal rate of HSCs or/and from a blockade or at least a slowdown of the LTHSC along their differentiation journey. It was also hypothesised that label-retaining HSCs (LR-HSCs), which divide minimally over time accumulate in old BM after completing four traceable symmetric self-renewal divisions to expand its size before entering a state of dormancy ^51^. Although, we could not directly address the question of self-renewal, we can argue based on our regulon and cell cycle analyses that old LTHSCs have kept their capacity to self-renew and did not reach a state of complete dormancy but reduced their proliferation linked to differentiation. Interestingly, we could associate this reduced and age-related proliferation/differentiation potential to a high level of Klf6 and Mycn activity, known to contribute to the stemness and self-renewal of different stem cells ^52^ and a high level of Gfi1b activity known to promote self-renewal of HSC ^53^.

One interesting outcome of our analysis is the link between the delay in differentiation and cell cycle activity changes of old HSPCs. First, we deduced from our computational cell cycle classification that lineage-primed HSPCs were less in G1/G0 than the LTHSC non-primed. This observation is fully consistent with current knowledge that the most undifferentiated HSCs reside in the G0 phase and cycle infrequently and that cell cycle overall becomes more frequent as HSCs are gradually committed ^40; 54^. Second, we detected an increase in HSPCs in G1/G0 phases in aged BM and old LTHSCs with increased in G1/G0 phases in comparison to young LTHSCs, reflecting the decrease in cell cycle activity of old HSCs when considered as a whole ^55^. Finally, when calculating a division rate per cells and studying division gene expression along the trajectory we could detect a loss of old dividing HSPCs located before the first bifurcation of the differentiation trajectory. These cells partially overlap in our trajectory with the div cluster, marked by genes related to asymmetric division such as *gpsm2*, *Ragcap* and *Ccnb1* ^56; 57^, suggesting that the delay in differentiation could be linked to an altered capacity of old HSPCs to divide asymmetrically. In addition, gene expression of cell cycle inhibitors clearly show that HSPCs at the beginning of the trajectory have increased expression in *Cdkn1a and Cdkn2c*, promoters of quiescence but a reduction in *Cdkn1b* activation, which promotes commitment ^58^. Interestingly, our analysis pointed out *Cdkn1a* as a direct target of Junb, and indirectly of Klf6. As the activation of *Cdkn1a* by Junb has been previously described to limit hematopoietic stem cell proliferation ^59^ and as Klf6 is a key factor in the Tgfbeta signalling pathway ^60; 61^, our work unveils an interesting pathway controlled by the cytokine Tgfbeta involving Klf6 as a key regulon and *Cdkn1a* as a cell cycle regulator that are enhanced upon aging and limit HSC differentiation.

In conclusion, our single-cell transcriptome-based identification of cell identity and its modifications associated with aging provides new information on cellular heterogeneity and intrinsic changes that will be useful for future investigation of the role of other regulators on the aged HSC phenotype.

## Acknowledgements

We thank Dr Lionel Spinelli for critically reading the manuscript and the code. We are grateful to the core flow cytometry and the animal facilities of the CRCM for providing supportive help and to the HaliodX and the TGML sequencing facilities for the single cell capture and sequencing. Computing resources for this study was provided both by the computing facilities DISC (Datacenter IT and Scientific Computing) of the Centre de Recherche en Cancérologie de Marseille and by the facilities of the Institut de Mathématiques de Marseille thanks to Dr Olivier Chabrol. This work was partly supported by the Excellence Initiative of Aix-Marseille University –A∗MIDEX, a French “Investissements d’Avenir” program to ED and ER), Institut Thématique Multi-Organisme-cancer, by l’Institut National du Cancer (grant number 20141PLBIO06 to ED) and the ARC foundation (PJA#20161204989 to ED). LH was the recipient of an interdisciplinary PhD grant from Aix Marseille University; MP’s postdoctoral fellowship was supported by the Fondation de France (#2017-00076284).

## MATERIALS AND METHODS

### Mouse model and cell sorting

C57BL/6-CD45.2 mice were purchased from Charles River Laboratories and were aged at the CRCM animal facility under specific pathogen-free conditions and according to the current European regulation. Only males were analysed, at young (2-3 months) and old (17-18 months) ages. HSPCs were collected from the BM of 5 young and 5 old mice over 2 independent batches with cells from 2 pooled young (Young_A sample) and 3 pooled old (Old_A sample) mice for one batch, and cells from 3 pooled young (Young_B sample) and 2 pooled old (Old_B sample) mice for the other one (Supplementary Table 1). For each sample, the BM was lineage depleted by using the Lineage Cell Depletion Kit (Miltenyi Biotec) and labelled with the following antibody cocktail: anti CD45.2, anti Sca-1, anti-cKit, anti CD150, anti Cd48, anti Cd34, and anti Flt3 antibodies (Supplementary Table 11) to purify Lin-Sca1+cKit+ Flt3-cells (HSPCs) by multi-parameter fluorescence-activated cell sorting (FACS) on a FACSAriaII (SpecialOrderResearch Products; BDBiosciences). Flow cytometry analyses were performed using a BD-LSRII cytometer and analysed using BD-DIVA Version 6.1.2 software (Special Order Research Products; BD Biosciences).

### Single cell RNA-seq and data processing

We used the 10x genomics platform from two facilities: HalioDX for samples Young_A and Old_A (Marseille, France) and TGML for samples Young_B and Old_B (Marseille, France). In both facilities, FACS purified HSPCs were loaded 30 min after the sorting onto a Chromium Single Cell Chip and processed with the Chromium Controller (10x Genomics) according to the manufacturer’s instructions for single cell barcoding at a target capture rate of 4000 individual cells per sample. Libraries were prepared using Chromium Single-Cell 3′ Reagent Kits v2 (10x Genomics) and were sequenced using an Illumina NextSeq500 sequencer to an average depth of about 45,000 reads per cell for Young_A and Old_B samples and about 130,000 reads per cell for Old_A and Young_B samples. Cell ranger software v2.2 was used to align reads to the (GRCm38) mm10 mouse reference genome. Cell counts and transcript detection rates are summarized in Supplementary Table 1.

### Quality control and data normalization

Cells outside 2 medians absolute deviation (MADs) from the median UMI log-counts were filtered out for each sample to discard poor quality cells and doublets. In total, 7433 young and 7482 old cells were kept. For each dataset (our four samples and the Rodriguez-Fraticelli dataset), genes with no UMI count in more than 0.5 percent of the cells were discarded. All gathering, 17513 genes were kept. Then, UMI counts were normalized with the NormalizeData Seurat function. For each cell, we considered the log transformation of the ratio of UMI counts per gene by the total UMI counts of the cell, multiply by a scaling factor of 10,000 (log(10,000(UMI_gene_/UMI_cell_)+1)).

### Cell cycle phase classification

Prediction of cell cycle phase for each cell was made with the cyclone ^62^, which relies on a pre-defined classifier for cell division constructed from a training dataset of synchronized mouse embryonic stem cells ^63^. For each cell a score based on raw count data before gene filtering was computed for each phase (G2/M, S and G1/G0) and used to assign a phase to the cell.

### HSPC subtype assignment

In order to assign known FACS cell identity in our HSPC pool, we used CaSTLe, a transfer learning method consisting in labeling cells in a scRNA-seq experiment, using knowledge learnt from other experiments on similar subtypes ^64^. We chose as source dataset a published scRNA-seq dataset obtained from FACS isolated HSPCs ^12^. Cells from this data set (approximately 2000 /per type) were divided into 4 subsets: the LTHSC (Lin- Sca1^+^ Kit^+^ Flt3^−^ Cd150^+^ Cd48^−^), the STHSC (Lin- Sca1^+^ Kit^+^ Flt3^−^ Cd150^−^ Cd48^−^), the MPP2s (Lin- Sca1^+^ Kit^+^ Flt3^−^ Cd150^+^ Cd48^+^), and the MPP3 (Lin- Sca1^+^ Kit^+^ Flt3^−^ Cd150^−^ Cd48^+^).

### Integration of the datasets

To minimize batch effect between datasets, we integrated our 4 sample datasets (Young_A, Young_B, Old_A, Old_B) following the procedure of Seurat 3 ^28^. Integration was done also for young and old conditions separately. Briefly, the highly variable genes for each dataset were selected with the FindVariableFeatures function (selection.method =“vst”) and ranked according to the number of datasets in which they were independently identified as highly variable. The top highly variable 2000 genes were thus integrated by iteratively merging pairs of datasets according to a given distance. Integration anchors, representing two cells that are predicted to originate from a common biological state in both datasets using a Canonical Correlation Analysis (CCA), was done using the FindIntegrationAnchors function (dims=1:15). Then, the expression of the target dataset was corrected using the difference in expression between the two expression vectors for each pair of anchor cells. This step was realized using IntegrateData function (dims= 1:15). This process resulted in an expression matrix that contains the batch-effect-corrected expression for the 2000 selected genes of the cells from the 4 samples.

### Data scaling and cell cycle regression

Standardised (*i.e.* centered and reduced) expression values with cell to cell variations due to cell cycle effect regressed were obtained with the ScaleData function of Seurat using the G2/M, S and G1/G0 scores previously computed for each cell by cyclone for the var.to.regress argument (cf Cell cycle phase classification).

### Dimension reduction and clustering

A PCA was performed on the scaled data using RunPCA Seurat function (npc = 40). The 15 first principal components of the PCA were kept for nonlinear dimension reduction and cell clustering. Uniform Manifold Approximation and Projection (UMAP), ^29^, a nonlinear dimension reduction method, was run using RunUMAP Seurat function package in order to embed cells in a 2-dimensional space. A K-nearest neighbour graph (KNN) based on the euclidean distance in PCA space was constructed (k = 20) to cluster the cells with the Louvain algorithm (resolution=0.5) using the FindNeighbors and FindClusters Seurat functions respectively.

### Pseudotime ordering

Unsupervised ordering of the HSPCs was done with the Seurat 3 integrated results as input to build a tree like differentiation trajectory using the DDRTree algorithm of the R package Monocle v2 ^32^. Integrated cells from: (i) all samples (young and old) excluding the primed pB cluster cells, (ii) young cells only or (iii) old cells only were processed with Monocle. For the three pseudotime ordering analyses (all cells, young only and old only), the 2000 gene expression matrix, scaled and regressed for cell cycle effect (see **Data scaling and cell cycle regression**) issued from the Seurat 3 integrated samples was loaded into Monocle using the newCellDataSet function (lowerDetectionLimit = 0.1, expressionFamily = uninormal()). The 2000 genes were set as ordering genes and trajectory building was made by calling the reduceDimension Monocle function (max_components = 2, reduction_method = ‘DDRTree’, norm_method = “none”, pseudo_expr = 0). For each of the three trajectories the root state was identified by selecting the Monocle state with the highest proportion of LTHSC predicted subtype (Fig. 4b; Supplementary Fig. 6B) in order to compute pseudotime values for the cells using the orderCells Monocle function. Expression of some genes as a function of pseudotime (Fig. 6g) was plotted with the plot_gene_expression Monocle function (using the Monocle normalization method with the estimateSizeFactor Monocle function).

### Differential gene expression analysis

Specific markers for each cluster (Supplementary Table 2) and for each Monocle state (Supplementary Table 8B) were identified using FindAllMarkers Seurat function, with default parameters. Genes significantly overexpressed in one cluster/state versus all the others (positive markers) were tested with Wilcoxon rank sum tests on the log-normalized data of the given cluster against all the others. DEGs between monocle state 2 and 4 (Supplementary Table 8C) were identified using the FindMarkers Seurat function. Only genes expressed in at least 10% of the cells in either of the two groups (min.pct = 0.1) and with a log fold change threshold of 0.25 (logfc.threshold = 0.25) were tested. A *p*-adjusted value (Bonferroni correction) threshold of 0.05 was applied to filter out non-significant markers.

Aging markers for the global population were obtained with the FindConservedMarkers Seurat function (min.pct = 0.1, logfc.threshold = 0) using the sequencing platform as grouping variable to minimize batch effect (Young_A, Old_A were processed on HalioDx platform and Young_B, Old_B on TGML platform). A Wilcoxon Rank sum test was performed on the log-normalized data between all young versus all old cells (Supplementary Table 5) from each batch separately and the two p-values for each gene were combined using the Tipett’s method. Genes presenting an opposite variation between the 2 batches were filtered out.

Aging markers for each cluster (Supplementary Table 6) and for each Monocle state (Supplementary Table 10B) were obtained with the same method by looking at the difference cluster per cluster and state per state (min.pct = 0.1, logfc.threshold = 0.25 for each cluster and min.pct = 0, logfc.threshold = 0 for each state). No tests were performed in the primed B cells clusters because it contained less than 3 cells in one young pool. From these results, for each cluster and each state only significant aging markers (combined p-value < 0.05 and same direction of variation in the 2 batches) were kept.

Among these markers the highly variable ones (average log fold change > 0.5 with aging in at least one cluster in both batches) were selected to generate heatmap for all clusters with primed clusters gathered (Fig. 3a) and for primed clusters only (Supplementary Fig. 4) with an adaptation of the DoHeatmap Seurat function with default parameters. Genes (raw) were ordered using hclust R function on standardised aging gene expression of the subset. Euclidian distance and unweighted pair group method with arithmetic mean (UPGMA) were used. Up and down regulated genes with aging were ordered separately.

Volcano plots for the global aging markers were drawn (Supplementary Fig. 3) with EnhancedVolcano function from the R package of the same name ^65^.

### Gene set enrichment analysis

To characterize the identified clusters with Seurat, we performed gene set enrichment analysis on cluster markers with g:Profiler v0.6.7 ^66^ with default arguments except for background set to all genes expressed in the whole dataset (i.e. genes that passed filtering during quality control). We tested enrichments in GO terms (GO:BP, GO:MF, GO:CC) as well as in terms from KEGG, REAC, TF, MI, CORUM, HP, HPA, OMIM databases (Fig. 1c and Supplementary Table 3). Cluster markers were also tested for enrichment in previously published gene set signatures related to HSPCs (Supplementary Table 12). Signatures tested were: Bcell_Chambers, Diff_Chambers, Gran_Chambers, HSC_Chambers, Lymph_Chambers, Mono_Chambers, Mye_Chambers, NK_Chambers, NaiveT_Chambers, and Ner_Chambers ^33^, lineage priming of HSC signatures C1, C2, C3, Mk, Er, Ba, Neu, Mo, Mo2, preDC, preB and preT ^12^, and HSCs and aging signatures Mm_HSC_Runx1_Wu, Mm_HSC_Tcf7_Wu ^67^, Mm_LT_HSC_Venezia, Mm_Proliferation_Venezia, Mm_Quiescence_Venezia ^68^, Polarity_factors_Ting and Novel_HSC_regul_polar_Ting ^57^. Cluster marker enrichment for the different signatures in comparaison to all dataset genes was tested using a hypergeometric test (phyper R function). To perform enrichment analysis of aging markers with a consistent gene number we gathered the overexpressed (resp. underexpressed) markers from at least one cluster and used gprofiler as describe above (Supplementary Table 7A & B). Expression scores of the signatures or of selected aging features from the enrichment analysis were calculated for each individual cell using the AddModuleScore Seurat function (on log-normalized data) with default parameters, using as input the genes of the signatures or the aging markers annotated for the selected features.

### Differential signature score analysis

Signature markers of Monocle state were tested in the same way as gene state markers (see above) using FindAllMarkers (min.pct= 0, logfc.threshold=0) with Student’s t-tests. Only signatures with an average score differences above 0.015 between one state versus all were kept. A *p*-adjusted value (Bonferroni correction) threshold of 0.05 was applied to filter out non-significant differences.

Signature score differences with aging in each state were tested in the same way as the aging markers per clusters (see above) using the FindConservedMarkers Seurat function (sequencing platform as grouping variable, min.pct and logfc.threshold set to 0) with Student’s t-tests. For each Monocle state, only average score differences of same sign and above 0.015 in the two batches presenting a combined p value < 0.05 were kept (Supplementary Table 8A).

The selected aging features expression score differences with aging in each cluster were tested in the same way as the aging markers per clusters (see above) using the FindConservedMarkers Seurat function (sequencing platform as grouping variable, min.pct and logfc.threshold set to 0) with Student’s t-tests (Supplementary Table 7C). For each cluster, only average score differences of same sign and above 0.1 in the two batches presenting a combined p value < 0.05 are considered as significant (Fig. 3b). No tests were performed in the primed B cells clusters because it contained less than 3 cells in one young pool.

### Regulons analysis

pySCENIC (1.10.0) was used with its command line implementation ^37^. The raw expression matrix for the cells of all samples was filtered, by keeping genes with a total expression greater than 2*0.01*(number of cell). 10698 genes passed the filtering. pyscenic grn command was used with grnboos2 method and default options and a fixed seed to derive co-expression modules between transcription factors and potential targets. We used as input all the markers of the Seurat clusters for which a transcription factor binding motif was available in the motifs-v9-nr.mgi-m0.001-o0.0 database provided by Scenic plus several TFs involved in early hematopoiesis, Spi1, Tal1, Zfpm1, Cbfa2t3, Erg, Fli1, Gata1, Gata2, Hhex, Runx1, Smad6 ^69^, Gfi1b ^70^, Zbtb16 ^71^. The obtained modules were refined by pruning targets that did not have an enrichment for a corresponding motif of the TF with pyscenic ctx command with –maskdropouts option using the motif database motifs-v9-nr.mgi-m0.001-o0.0 and the cis-target database mm9-tss-centered-10kb-7species.mc9nr. Only positive regulons (i.e. those with a positive correlation between the TF and its targets) were kept for downstream analysis (Supplementary Table 9A). AUCell scores (regulon activities) in each cell were computed with pycenic aucell command (default options). To be noted that number of target genes was highly variable from a regulon to another (Supplementary Fig. 9)

For young and old HSPCs, two Heatmaps of regulon activity scores, along pseudotime were made, in order to analyze transcriptional activity at the two bifurcation points for both ages. See supplemental methods for detailed regulons heatmaps construction.

Regulon markers of monocle states were tested in the same way as gene state markers (see above) with their AUCell scores using FindAllMarkers Seurat function (min.pct= 0.1, logfc.threshold=0) with Wilcoxon rank sum tests. Only regulon with an average AUCell score differences above 0.002 between one state versus all the others were kept. A *p*-adjusted value (Bonferroni correction) threshold of 0.05 was applied to filter out non-significant differences.

Regulon activity differences with aging in each state were tested in the same way as the aging markers per clusters (see above) using the FindConservedMarkers Seurat function (sequencing platform as grouping variable, min.pct = 0.1 and logfc.threshold = 0) with Wilcoxon rank sum tests. For each state, only average AUCell score differences of same sign and above 0.002 in the two batches presenting a combined p value < 0.05 were kept (Supplementary Table 9B).

### Analysis of HSPC subtypes and cell cycle phases in the differentiation trajectory depending on age

Cell Density (Fig. 6d), division rate (Fig. 6e) and stacked plot of HSPC subtypes (Fig. 6f) were computed and plotted along pseudotime at each age for the 3 HSPC fates: toward the T lymphocyte priming (Monocle states 1 and 2), toward the Mastocytes/Neutrophils priming (Monocle states 1, 3 and 4) and toward the Megakaryocytes/Erythrocytes priming Monocle states 1, 3 and 5). For division rate and stacked plot of HSPC subtypes, pseudotime was cut into 50 bins. For each age, in each pseudotime bin, division rate was computed as the ratio of the number of cells with a G2/M phase assigned to the total number of cells in the bin.

### Statistics

Statistics were computed with R software v3.5.1. The statistical tests for gene expression and signature or regulon activity scores were performed with Seurat and are detailed above. In each cluster and in non-primed/primed clusters gathered, the enrichment of age was tested using a hypergeometric test (phyper R function Fig. 2b). Chi2 tests (chisq.test R function) were performed to test independence between cell cycle phase and age, in all cells (Fig. 6a) and in each HSPC subtype separately (Fig. 6b), and in the cells at the departure of the trajectory (Pseudotime <2, Supplementary Fig. 10) and to test independence between Monocle state and age in all Monocle states (Fig. 4i) and in the states 2, 3 and 5 only (Fig. 4j). Fisher’s exact test (fisher.test R function) were performed to test independence between Monocle state and age in each Seurat cluster (Supplementary Fig. 8B). Wilcoxon rank sum test was used to test for median difference between pseudotime value distributions of young and old cells (Fig. 4h). In each cluster a linear regression was computed between the average log fold change (in the cluster) and the global (in all cells) average log fold change of the aging markers recovered in the cluster (lm R function Supplementary Fig. 5). Smooth curves of module score expression in pseudotime through the different fates for young and old cells were drawn for quiescence and proliferation signature (Fig. 6C) using the geom_smooth function ggplot2 R package ^72^ with the gam function of mgcv R package ^73^.

### Data and code availability

The single-cell RNA-seq data generated here are available in the Gene Expression Omnibus database under accession code GSE147729. All R and python codes used for data analysis are integrated in a global snakemake workflow available at: https://gitcrcm.marseille.inserm.fr/herault/scHSC_herault

## References

1. Spangrude GJ, Heimfeld S, Weissman IL. Purification and characterization of mouse hematopoietic stem cells. Science 1988, 241(4861): 58–62.

2. Osawa M, Hanada K, Hamada H, Nakauchi H. Long-term lymphohematopoietic reconstitution by a single CD34-low/negative hematopoietic stem cell. Science 1996, 273(5272): 242–245.

3. Akashi K, Traver D, Miyamoto T, Weissman IL. A clonogenic common myeloid progenitor that gives rise to all myeloid lineages. Nature 2000, 404(6774): 193–197.

4. Adolfsson J, Mansson R, Buza-Vidas N, Hultquist A, Liuba K, Jensen CT, et al. Identification of Flt3+ lympho-myeloid stem cells lacking erythro-megakaryocytic potential a revised road map for adult blood lineage commitment. Cell 2005, 121(2): 295–306.

5. Kiel MJ, Yilmaz OH, Iwashita T, Yilmaz OH, Terhorst C, Morrison SJ. SLAM family receptors distinguish hematopoietic stem and progenitor cells and reveal endothelial niches for stem cells. Cell 2005, 121(7): 1109–1121.

6. Oguro H, Ding L, Morrison SJ. SLAM family markers resolve functionally distinct subpopulations of hematopoietic stem cells and multipotent progenitors. Cell Stem Cell 2013, 13(1): 102–116.

7. Dykstra B, Olthof S, Schreuder J, Ritsema M, de Haan G. Clonal analysis reveals multiple functional defects of aged murine hematopoietic stem cells. J Exp Med 2011, 208(13): 2691–2703.

8. Morita Y, Ema H, Nakauchi H. Heterogeneity and hierarchy within the most primitive hematopoietic stem cell compartment. J Exp Med 2010, 207(6): 1173–1182.

9. Yamamoto R, Morita Y, Ooehara J, Hamanaka S, Onodera M, Rudolph KL, et al. Clonal analysis unveils self-renewing lineage-restricted progenitors generated directly from hematopoietic stem cells. Cell 2013, 154(5): 1112–1126.

10. Naik SH, Perie L, Swart E, Gerlach C, van Rooij N, de Boer RJ, et al. Diverse and heritable lineage imprinting of early haematopoietic progenitors. Nature 2013, 496(7444): 229–232.

11. Paul F, Arkin Y, Giladi A, Jaitin DA, Kenigsberg E, Keren-Shaul H, et al. Transcriptional Heterogeneity and Lineage Commitment in Myeloid Progenitors. Cell 2015, 163(7): 1663–1677.

12. Rodriguez-Fraticelli AE, Wolock SL, Weinreb CS, Panero R, Patel SH, Jankovic M, et al. Clonal analysis of lineage fate in native haematopoiesis. Nature 2018, 553(7687): 212–216.

13. Haas S, Trumpp A, Milsom MD. Causes and Consequences of Hematopoietic Stem Cell Heterogeneity. Cell Stem Cell 2018, 22(5): 627–638.

14. Zhang Y, Gao S, Xia J, Liu F. Hematopoietic Hierarchy - An Updated Roadmap. Trends Cell Biol 2018, 28(12): 976–986.

15. Laurenti E, Gottgens B. From haematopoietic stem cells to complex differentiation landscapes. Nature 2018, 553(7689): 418–426.

16. Baldridge MT, King KY, Boles NC, Weksberg DC, Goodell MA. Quiescent haematopoietic stem cells are activated by IFN-gamma in response to chronic infection. Nature 2010, 465(7299): 793–797.

17. Haas S, Hansson J, Klimmeck D, Loeffler D, Velten L, Uckelmann H, et al. Inflammation-Induced Emergency Megakaryopoiesis Driven by Hematopoietic Stem Cell-like Megakaryocyte Progenitors. Cell Stem Cell 2015, 17(4): 422–434.

18. Geiger H, de Haan G, Florian MC. The ageing haematopoietic stem cell compartment. Nat Rev Immunol 2013, 13(5): 376–389.

19. Chung SS, Park CY. Aging, hematopoiesis, and the myelodysplastic syndromes. Blood Adv 2017, 1(26): 2572–2578.

20. Sun D, Luo M, Jeong M, Rodriguez B, Xia Z, Hannah R, et al. Epigenomic profiling of young and aged HSCs reveals concerted changes during aging that reinforce self-renewal. Cell Stem Cell 2014, 14(5): 673–688.

21. Li X, Zeng X, Xu Y, Wang B, Zhao Y, Lai X, et al. Mechanisms and rejuvenation strategies for aged hematopoietic stem cells. J Hematol Oncol 2020, 13(1): 31.

22. Cooper JN, Young NS. Clonality in context: hematopoietic clones in their marrow environment. Blood 2017, 130(22): 2363–2372.

23. Beerman I, Bhattacharya D, Zandi S, Sigvardsson M, Weissman IL, Bryder D, et al. Functionally distinct hematopoietic stem cells modulate hematopoietic lineage potential during aging by a mechanism of clonal expansion. Proc Natl Acad Sci U S A 2010, 107(12): 5465–5470.

24. Yamamoto R, Wilkinson AC, Ooehara J, Lan X, Lai CY, Nakauchi Y, et al. Large-Scale Clonal Analysis Resolves Aging of the Mouse Hematopoietic Stem Cell Compartment. Cell Stem Cell 2018, 22(4): 600–607 e604.

25. de Haan G, Lazare SS. Aging of hematopoietic stem cells. Blood 2018, 131(5): 479–487.

26. Grover A, Sanjuan-Pla A, Thongjuea S, Carrelha J, Giustacchini A, Gambardella A, et al. Single-cell RNA sequencing reveals molecular and functional platelet bias of aged haematopoietic stem cells. Nat Commun 2016, 7:11075.

27. Kowalczyk MS, Tirosh I, Heckl D, Rao TN, Dixit A, Haas BJ, et al. Single-cell RNA-seq reveals changes in cell cycle and differentiation programs upon aging of hematopoietic stem cells. Genome Res 2015, 25(12): 1860–1872.

28. Stuart T, Butler A, Hoffman P, Hafemeister C, Papalexi E, Mauck WM, 3rd, et al. Comprehensive Integration of Single-Cell Data. Cell 2019, 177(7): 1888–1902 e1821.

29. McInnes L, Healy J, Melville J. UMAP: uniform manifold approximation and projection for dimension reduction. Preprint at https://arxiv.org/abs/1802.03426; 2018.

30. Sanjuan-Pla A, Macaulay IC, Jensen CT, Woll PS, Luis TC, Mead A, et al. Platelet-biased stem cells reside at the apex of the haematopoietic stem-cell hierarchy. Nature 2013, 502(7470): 232–236.

31. Zidi B, Vincent-Fabert C, Pouyet L, Seillier M, Vandevelde A, N’Guessan P, et al. TP53INP1 deficiency maintains murine B lymphopoiesis in aged bone marrow through redox-controlled IL-7R/STAT5 signaling. Proc Natl Acad Sci U S A 2019, 116(1): 211–216.

32. Qiu X, Mao Q, Tang Y, Wang L, Chawla R, Pliner HA, et al. Reversed graph embedding resolves complex single-cell trajectories. Nat Methods 2017, 14(10): 979–982.

33. Chambers SM, Boles NC, Lin KY, Tierney MP, Bowman TV, Bradfute SB, et al. Hematopoietic fingerprints: an expression database of stem cells and their progeny. Cell Stem Cell 2007, 1(5): 578–591.

34. Chen X, Deng H, Churchill MJ, Luchsinger LL, Du X, Chu TH, et al. Bone Marrow Myeloid Cells Regulate Myeloid-Biased Hematopoietic Stem Cells via a Histamine-Dependent Feedback Loop. Cell Stem Cell 2017, 21(6): 747–760 e747.

35. Lu YC, Sanada C, Xavier-Ferrucio J, Wang L, Zhang PX, Grimes HL, et al. The Molecular Signature of Megakaryocyte-Erythroid Progenitors Reveals a Role for the Cell Cycle in Fate Specification. Cell Rep 2018, 25(8): 2083–2093 e2084.

36. Muench DE, Grimes HL. Transcriptional Control of Stem and Progenitor Potential. Curr Stem Cell Rep 2015, 1(3): 139–150.

37. Aibar S, Gonzalez-Blas CB, Moerman T, Huynh-Thu VA, Imrichova H, Hulselmans G, et al. SCENIC: single-cell regulatory network inference and clustering. Nat Methods 2017, 14(11): 1083–1086.

38. Moroy T, Vassen L, Wilkes B, Khandanpour C. From cytopenia to leukemia: the role of Gfi1 and Gfi1b in blood formation. Blood 2015, 126(24): 2561–2569.

39. Pietras EM, Warr MR, Passegue E. Cell cycle regulation in hematopoietic stem cells. J Cell Biol 2011, 195(5): 709–720.

40. Seita J, Weissman IL. Hematopoietic stem cell: self-renewal versus differentiation. Wiley Interdiscip Rev Syst Biol Med 2010, 2(6): 640–653.

41. Orford KW, Scadden DT. Deconstructing stem cell self-renewal: genetic insights into cell-cycle regulation. Nat Rev Genet 2008, 9(2): 115–128.

42. Velten L, Haas SF, Raffel S, Blaszkiewicz S, Islam S, Hennig BP, et al. Human haematopoietic stem cell lineage commitment is a continuous process. Nat Cell Biol 2017, 19(4): 271–281.

43. Notta F, Zandi S, Takayama N, Dobson S, Gan OI, Wilson G, et al. Distinct routes of lineage development reshape the human blood hierarchy across ontogeny. Science 2016, 351(6269): aab2116.

44. Notta F, Doulatov S, Laurenti E, Poeppl A, Jurisica I, Dick JE. Isolation of single human hematopoietic stem cells capable of long-term multilineage engraftment. Science 2011, 333(6039): 218–221.

45. Pietras EM, Reynaud D, Kang YA, Carlin D, Calero-Nieto FJ, Leavitt AD, et al. Functionally Distinct Subsets of Lineage-Biased Multipotent Progenitors Control Blood Production in Normal and Regenerative Conditions. Cell Stem Cell 2015, 17(1): 35–46.

46. Carrelha J, Meng Y, Kettyle LM, Luis TC, Norfo R, Alcolea V, et al. Hierarchically related lineage-restricted fates of multipotent haematopoietic stem cells. Nature 2018, 554(7690): 106–111.

47. Benayoun BA, Pollina EA, Singh PP, Mahmoudi S, Harel I, Casey KM, et al. Remodeling of epigenome and transcriptome landscapes with aging in mice reveals widespread induction of inflammatory responses. Genome Res 2019, 29(4): 697–709.

48. Xia S, Zhang X, Zheng S, Khanabdali R, Kalionis B, Wu J, et al. An Update on Inflamm-Aging: Mechanisms, Prevention, and Treatment. J Immunol Res 2016, 2016: 8426874.

49. Challen GA, Boles NC, Chambers SM, Goodell MA. Distinct hematopoietic stem cell subtypes are differentially regulated by TGF-beta1. Cell Stem Cell 2010, 6(3): 265–278.

50. Klimmeck D, Cabezas-Wallscheid N, Reyes A, von Paleske L, Renders S, Hansson J, et al. Transcriptome-wide profiling and posttranscriptional analysis of hematopoietic stem/progenitor cell differentiation toward myeloid commitment. Stem Cell Reports 2014, 3(5): 858–875.

51. Bernitz JM, Kim HS, MacArthur B, Sieburg H, Moore K. Hematopoietic Stem Cells Count and Remember Self-Renewal Divisions. Cell 2016, 167(5): 1296–1309 e1210.

52. Park CS, Lewis A, Chen T, Lacorazza D. Concise Review: Regulation of Self-Renewal in Normal and Malignant Hematopoietic Stem Cells by Kruppel-Like Factor 4. Stem Cells Transl Med 2019, 8(6): 568–574.

53. Zeng H, Yucel R, Kosan C, Klein-Hitpass L, Moroy T. Transcription factor Gfi1 regulates self-renewal and engraftment of hematopoietic stem cells. EMBO J 2004, 23(20): 4116–4125.

54. Cheung TH, Rando TA. Molecular regulation of stem cell quiescence. Nat Rev Mol Cell Biol 2013, 14(6): 329–340.

55. Flach J, Bakker ST, Mohrin M, Conroy PC, Pietras EM, Reynaud D, et al. Replication stress is a potent driver of functional decline in ageing haematopoietic stem cells. Nature 2014, 512(7513): 198–202.

56. Zhu J, Wen W, Zheng Z, Shang Y, Wei Z, Xiao Z, et al. LGN/mInsc and LGN/NuMA complex structures suggest distinct functions in asymmetric cell division for the Par3/mInsc/LGN and Galphai/LGN/NuMA pathways. Mol Cell 2011, 43(3): 418–431.

57. Ting SB, Deneault E, Hope K, Cellot S, Chagraoui J, Mayotte N, et al. Asymmetric segregation and self-renewal of hematopoietic stem and progenitor cells with endocytic Ap2a2. Blood 2012, 119(11): 2510–2522.

58. Hao S, Chen C, Cheng T. Cell cycle regulation of hematopoietic stem or progenitor cells. Int J Hematol 2016, 103(5): 487–497.

59. Santaguida M, Schepers K, King B, Sabnis AJ, Forsberg EC, Attema JL, et al. JunB protects against myeloid malignancies by limiting hematopoietic stem cell proliferation and differentiation without affecting self-renewal. Cancer Cell 2009, 15(4): 341–352.

60. Botella LM, Sanz-Rodriguez F, Komi Y, Fernandez LA, Varela E, Garrido-Martin EM, et al. TGF-beta regulates the expression of transcription factor KLF6 and its splice variants and promotes co-operative transactivation of common target genes through a Smad3-Sp1-KLF6 interaction. Biochem J 2009, 419(2): 485–495.

61. Dhaouadi N, Li JY, Feugier P, Gustin MP, Dab H, Kacem K, et al. Computational identification of potential transcriptional regulators of TGF-ss1 in human atherosclerotic arteries. Genomics 2014, 103(5-6): 357–370.

62. Scialdone A, Natarajan KN, Saraiva LR, Proserpio V, Teichmann SA, Stegle O, et al. Computational assignment of cell-cycle stage from single-cell transcriptome data. Methods 2015, 85:54–61.

63. Buettner F, Natarajan KN, Casale FP, Proserpio V, Scialdone A, Theis FJ, et al. Computational analysis of cell-to-cell heterogeneity in single-cell RNA-sequencing data reveals hidden subpopulations of cells. Nat Biotechnol 2015, 33(2): 155–160.

64. Lieberman Y, Rokach L, Shay T. CaSTLe - Classification of single cells by transfer learning: Harnessing the power of publicly available single cell RNA sequencing experiments to annotate new experiments. PLoS One 2018, 13(10): e0205499.

65. Blighe K, Rana S, Lewis M. EnhancedVolcano: Publication-ready volcano plots with enhanced colouring and labeling. 2018.

66. Reimand J, Kull M, Peterson H, Hansen J, Vilo J. g:Profiler-a web-based toolset for functional profiling of gene lists from large-scale experiments. Nucleic Acids Res 2007, 35(Web Server issue): W193–200.

67. Wu JQ, Seay M, Schulz VP, Hariharan M, Tuck D, Lian J, et al. Tcf7 is an important regulator of the switch of self-renewal and differentiation in a multipotential hematopoietic cell line. PLoS Genet 2012, 8(3): e1002565.

68. Venezia TA, Merchant AA, Ramos CA, Whitehouse NL, Young AS, Shaw CA, et al. Molecular signatures of proliferation and quiescence in hematopoietic stem cells. PLoS Biol 2004, 2(10): e301.

69. Bonzanni N, Garg A, Feenstra KA, Schutte J, Kinston S, Miranda-Saavedra D, et al. Hard-wired heterogeneity in blood stem cells revealed using a dynamic regulatory network model. Bioinformatics 2013, 29(13): i80–88.

70. Schutte J, Wang H, Antoniou S, Jarratt A, Wilson NK, Riepsaame J, et al. An experimentally validated network of nine haematopoietic transcription factors reveals mechanisms of cell state stability. Elife 2016, 5:e11469.

71. Poplineau M, Vernerey J, Platet N, N’Guyen L, Herault L, Esposito M, et al. PLZF limits enhancer activity during hematopoietic progenitor aging. Nucleic Acids Res 2019, 47(9): 4509–4520.

72. Wickham H. ggplot2: Elegant Graphics for Data Analysis. Springer-Verlag New York 2016.

73. Wood SN, Pya N, Säfken B. Smoothing Parameter and Model Selection for General Smooth Models. Journal of the American Statistical Association 2016, 111(516): 1548–1563.

